# Genetically-encoded phase separation sensors for intracellular probing of biomolecular condensates

**DOI:** 10.1101/2024.08.29.610365

**Authors:** Alexa Regina Chua Avecilla, Jeremy Thomas, Felipe Garcia Quiroz

**Affiliations:** Wallace H. Coulter Department of Biomedical Engineering, Georgia Institute of Technology and Emory University, Atlanta, Georgia, USA

**Author notes:** **Correspondence**: Felipe Garcia Quiroz, Wallace H. Coulter Department of Biomedical Engineering, Emory University, Atlanta, Georgia 30322, USA.

## Abstract

Biomolecular condensates are dynamic membraneless compartments with enigmatic roles across intracellular phenomena. Intrinsically-disordered proteins (IDPs) often function as condensate scaffolds, fueled by their liquid-liquid phase separation (LLPS) dynamics. Intracellular probing of these condensates relies on live-cell imaging of IDP-scaffolds tagged with fluorescent proteins. Conformational heterogeneity in IDPs, however, renders them uniquely sensitive to molecular-level fusions, risking distortion of the native biophysical properties of IDP-scaffolds and their assemblies. Probing epidermal condensates in mouse skin, we recently introduced genetically encoded LLPS-sensors that circumvent the need for molecular-level tagging of skin IDPs. The concept of LLPS-sensors involves a shift in focus from subcellular tracking of IDP-scaffolds to higher-level observations that report on the assembly and liquid-dynamics of their condensates. Towards advancing the repertoire of intracellular LLPS-sensors, here we demonstrate biomolecular approaches for the evolution and tunability of epidermal LLPS-sensors and assess their impact in early and late stages of intracellular LLPS dynamics. Benchmarking against scaffold-bound fluorescent reporters, we found that tunable ultraweak scaffold-sensor interactions are key to the sensitive and innocuous probing of nascent and established biomolecular condensates. Our LLPS-sensitive tools pave the way for the high-fidelity intracellular probing of IDP-governed biomolecular condensates across biological systems.

## Introduction

Biomolecular condensates recently emerged as ubiquitous intracellular membraneless compartments with enigmatic liquid-like properties^1–5^. These incompletely understood condensates orchestrate a wide range of vital molecular mechanisms at cell and tissue levels ^6–9^. In addition to roles in homeostasis and stress response, aberrant biomolecular condensates are implicated in neurodegenerative diseases^5,10–12^ and cancer^12–14^. Intrinsically disordered proteins (IDPs) and large intrinsically disordered regions (IDRs) underpin the formation and liquid-like dynamics of numerous intracellular condensates^1,4,15,16^. They function as condensate “scaffolds”, driven by a concentration-dependent process of liquid-liquid phase separation (LLPS) or related phase transitions^17^. Above a specific concentration unique to the IDP-scaffold, known as the saturation concentration (C_sat_), they segregate into two distinct phases: a dense phase of IDP-rich droplets and a dilute phase of dispersed IDP-scaffolds^2,4^. In addition to defining condensate self-assembly dynamics, IDP-scaffolds may recruit “client” biomolecules through specific binding domains or weak IDP-IDP interactions^3,18,19^, expanding condensate composition and functionality.^20,21^

The underlying LLPS dynamics are sensitive to environmental and molecular-level perturbations. Two salient examples are post-translational modifications of IDP-scaffolds and physiological fluctuations in pH and ions, which dramatically alter condensate assembly dynamics and material properties^22,23^. Probing these functional dynamics intracellularly requires live-cell methods. State-of-the-art approaches involve the tagging of IDP-scaffolds with fluorescent proteins for live-cell imaging, or with enzymes for proximity-dependent biotinylation and proteomics ^3,4^. Live imaging probes the biophysical properties of intracellular condensates, while proximity proteomics enables in situ and time-dependent analyses of their biomolecular composition ^24,25^.

Challenging these live-cell efforts, the molecular-level tagging of native IDP-scaffolds risks unpredictably altering their intracellular localization and underlying LLPS dynamics^7,26^. This is in part because IDP-scaffolds functionally exploit high-entropy conformational dynamics that are sensitive to their molecular neighborhood ^27–29^. For neurodegeneration-relevant IDP assemblies, tagging with fluorescent proteins for live imaging has been shown to alter their composition, ultrastructure and toxicity ^30–32^. Furthermore, IDP-scaffolds may be regulated in vivo through proteolysis, limiting the utility of N-terminal or C-terminal protein tags. Filaggrin (FLG) exemplifies this regulation, undergoing N-terminal processing in the skin soon after condensate assembly ^7^. Through the lens of bioengineering, the immunogenicity of fluorescent protein tags ^33,34^ threatens clinical translation of synthetic IDP-scaffolds and synthetic condensates^35^ in human cells.

We currently lack tools for high-fidelity live-cell probing of intracellular condensates without scaffold-level tagging. Emerging approaches such as label-free microscopy coupled with deep learning appear promising for the identification of solid-like neuropathological IDP aggregates in cell monolayers ^32^. We suspect that related approaches may also succeed with a subset of IDP-driven condensates in cultured cells. Even with sustained progress in label-free methods, biomolecular tools will be needed to achieve high-resolution and innocuous probing of condensate dynamics at biophysical and biochemical levels.

In beginning to address this challenge, we recently introduced fluorescent IDP-based LLPS- sensors that are sensitive to the ultra-weak intermolecular interactions that drive the formation and LLPS dynamics of epidermal condensates^7^. We showed that live-cell probing of sensor signal within epidermal condensates in keratinocytes mirrored expected changes in their liquid- dynamics. In mice genetically engineered to express these epidermal LLPS-sensors and other subcellular markers, we unearthed previously enigmatic keratohyalin granules (KGs) as liquid-like epidermal condensates whose assembly and pH-triggered disassembly drive skin barrier formation^7^.

To stimulate the development of novel genetically encoded LLPS-sensors, here we dissect the biomolecular engineering and features of epidermal LLPS-sensors at two key levels: (1) their sensitive marking of biomolecular condensates (i.e., high signal-to-noise ratio for live-cell approaches), and (2) the impact of LLPS-sensors on early and late stages of intracellular phase separation. Excitingly, our data demonstrate two notable properties of LLPS-sensors: (i) highly tunable range of sensitivity, and (ii) innocuous probing of intracellular LLPS dynamics. Benchmarking our top-performing LLPS-sensor against a scaffold-bound fluorescent reporter- client, we discovered that ultraweak scaffold-sensor interactions are key to the high-fidelity probing of nascent and established biomolecular condensates. Our approaches and tools demonstrate a path toward rigorous intracellular probing of IDP-governed biomolecular condensates across biological systems.

## Results and discussion

### Concept and evolution of LLPS-sensors

The concept of LLPS-sensors involves a shift in focus from subcellular tracking of IDP-scaffolds to higher-level observations that report on the assembly and liquid-dynamics of their condensates. This shift eliminates the need to directly label the IDP-scaffold, creating the opportunity to engineer a protein that is sensitive to the underlying density phase transition (Fig. 1A-B). We hypothesized that an engineered IDP that shares LLPS-relevant information (e.g., sequences responsible for charge-charge, cation-pi, pi-pi, hydrogen bonding, and hydrophobic interactions) with an IDP-scaffold can be evolved to engage in weak but multivalent interactions that only become substantial when the IDP-scaffold resides within dense condensates. If fused to a fluorescent protein, the resulting LLPS-sensor would experience a dramatic gain in signal-to-noise ratio as it accumulates within nascent condensates (Fig. 1A).

**Fig 1.**
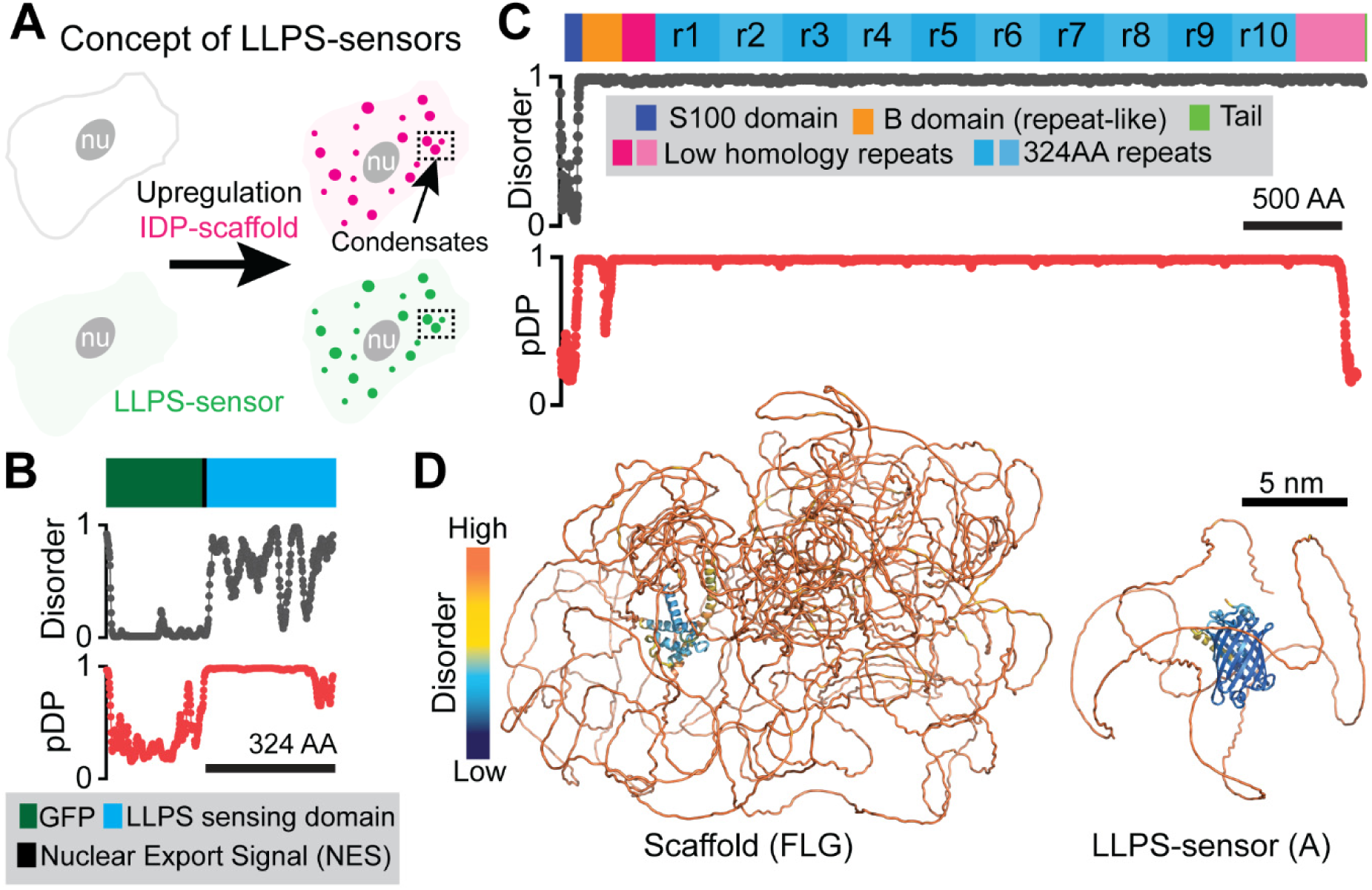
Concept and molecular features of genetically encoded LLPS-sensors. **(A)** Exploiting IDP-encoded LLPS information from a target IDP-scaffold, a fluorescent LLPS-sensor switches from a diffuse (dim green; left) state to a condensate-enriched (bright; right) state as intracellular upregulation of an IDP-scaffold results in LLPS-driven assembly of biomolecular condensates (magenta). Without artifacts from molecular-level fluorescent tagging of IDP-scaffolds, live-cell imaging of LLPS-sensor fluorescence reveals the native intracellular LLPS behavior of (untagged, non-fluorescent) IDP-scaffolds and their condensates. **(B)** Domain architecture, and corresponding disorder (where 1=disorder) and droplet-promoting propensity (pDP; where 1 indicates highest LLPS information content) plots for a recently developed LLPS-sensor (Sensor A) deployed to probe cytoplasmic epidermal condensates in mice^7^. **(C)** Domain architecture, and corresponding disorder and droplet-promoting propensity plots for FLG, the His-rich IDP-scaffold of human epidermal condensates. **(D)** AlphaFold2 predictions of the molecular 3D structure and disorder for FLG and Sensor A.

We previously reported that Sensor A (Fig. 1B), an engineered IDP that incorporates the LLPS-relevant information of FLG (Fig. 1C), successfully exposed FLG-driven intracellular LLPS dynamics in the skin of genetically engineered mice. We also showed that Sensor A could probe untagged endogenous epidermal condensates in mouse and human skin, despite mouse FLG and human FLG lacking sequence conservation. Sensor A is comprised of two domains: (i) an engineered IDP with little sequence identity to FLG repeats, and (2) an engineered green fluorescent protein (Fig.1B, 1D). FLG exhibits high disorder and LLPS-relevant information across nearly the entirety of its repetitive architecture (Fig. 1C-D). Here we take advantage of this repetitive architecture to probe FLG-like (r8)_n_ IDP-scaffolds with variable (n) copies of the domain r8 (Fig. S1A-B), creating a testbed to study the evolution and performance of LLPS-sensors.

To examine the tunability of epidermal LLPS-sensors, we treated mRFP1-(r8)_8_-Ctail scaffolds (Fig. S1A) and their condensates as mimics of FLG and their KGs. In the case of FLG and notably mRFP1-(r8)_8_-Ctail, a single repeat domain (r8) contains all the relevant LLPS-information (Fig. 1C and Fig. S1B). Importantly, we do not interpret the high droplet-promoting scores in Fig. 1B-C as reflective of a high probability of intracellular phase separation. We treat them as a metric of LLPS- information content, since we previously showed that (r8)_1_ and (r8)_2_ do not drive intracellular condensate formation even at extremely high expression levels^7^. Sensor A, as a reference, uses an IDP-sensing domain (ir8H2) that shares little sequence identity to r8 (21.4%) but preserves the overall amino acid composition of r8^7^ and hence its high droplet-promoting scores (Fig. 1B and Fig. S1B).

We co-transfected genes encoding a library of LLPS-sensor designs (Table S1) and mRFP1- (r8)_8_-Ctail into immortalized human keratinocytes (HaCATs) under conditions where no endogenous FLG is expressed. We note that FLG is a marker of late epidermal differentiation and stratification. Focusing on the IDP-sensing domain, we considered r8 as the simplest IDP that shares LLPS-information with our scaffold (Fig. 2A and Fig. S1C). We fused r8 to a nuclear export signal (LELLEDLTL)^36^ and to superfolder GFP (sfGFP; Table S2) with the same architecture as Sensor A in Fig. 1B. Despite the r8 domain being shared between this putative LLPS-sensor and the target IDP-scaffold, this initial sensor design appeared predominantly diffuse throughout the cytosol with little sensor signal co-localized exclusively with mRFP1-(r8)_8_-Ctail condensates (Fig. 2B). Quantifying their recruitment to condensates, we measured the partition coefficient (*P*), that is the ratio of LLPS-sensor signal present within condensates compared to the cytosol. The sfGFP-tagged r8 design exhibited a low average partition coefficient (Fig. 2C; *P*= 2.13 ± 0.55). The poor performance of this primitive sensor design showed the need to evolve LLPS-sensors for sensitive marking of target condensates.

**Fig 2:**
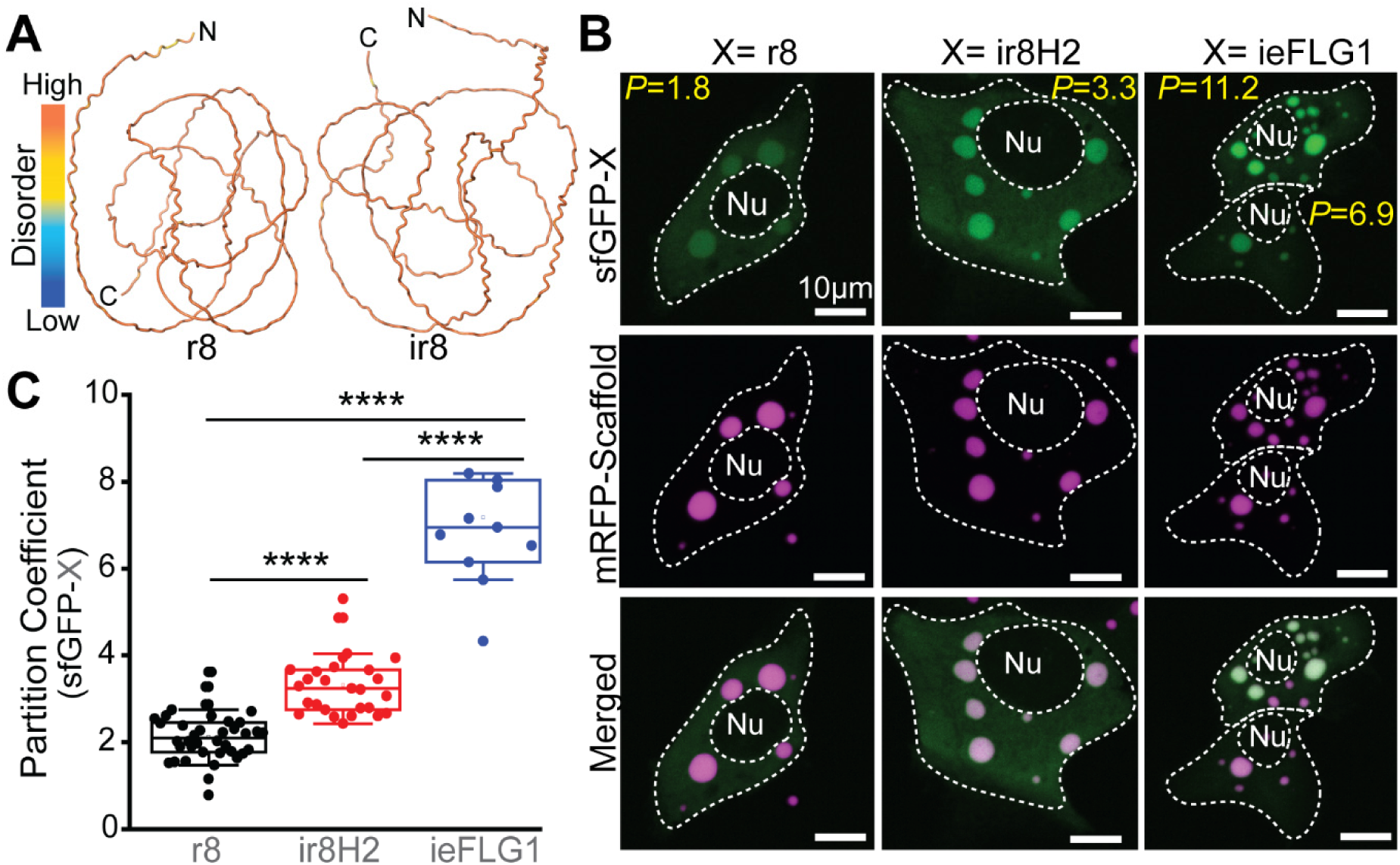
Sequence heuristics to encode LLPS in IDPs guide the evolution of tunable LLPS-sensing domains. **(A)** AlphaFold2 prediction of the 3D structure and disorder for a native FLG repeat domain (r8; Fig. 1C) and a dissimilar domain (ir8) generated by reversing the sequence of r8 —willfully writing the r8 sequence from C-terminus to N-terminus. This sequence reversal destroys sequence similarity, while conserving intrinsic disorder and overall amino acid composition, which helps preserve key LLPS information in IDPs. The r8 domain encodes LLPS-relevant information (Fig. S1A-B) but does not drive intracellular LLPS ^7^. **(B-C)** Live-cell images (**B**) and quantifications (**C**) of LLPS-sensor variants partitioning into mRFP1-(r8)_8_-Ctail condensates. The variants share a fixed fluorescent protein domain (sfGFP) but differ in the LLPS-sensing domain (X). ir8H2: variant of ir8 with H>Y mutations. ieFLG1: a de novo engineered r8-like sequence enriched for LLPS-relevant features. *P*: Partition Coefficient. Nu: nucleus. **(C)** Each data point corresponds to the LLPS-sensor partition coefficient across all sizable condensates in a cell. Asterisks denote statistical significance (p≤0.0001).

To optimize the IDP-sensing domain we turned to IDPs with low sequence identity to r8. In principle, sustained intracellular expression of LLPS-sensors with high sequence identity to IDP- scaffolds risks disruption of functional scaffold-specific biomolecular interactions. We applied a simple but dramatic sequence modification to r8: inversion of the syntax by reading its sequence in the unorthodox C- to N-terminal direction. The resulting protein (ir8) minimizes sequence identity to r8 (25.6%) but preserves its overall composition and disorder (Fig. 2A). Unlike scrambling, this strategy keeps the relative spacing and patterning of residues that function as stickers and spacers, which are important sources of LLPS-information ^37–39^. We note, however, that sequence inversion of IDP domains can shift LLPS dynamics^7,40,41^.

Using the ir8 domain, we introduced 18 histidine (His) to tyrosine (Tyr) mutations that mapped to naturally-occurring His>Tyr mutations in r8 and r9 of FLG — accounting for ∼50% of the 37 His residues in r8. These His>Tyr mutations are known to increase LLPS propensity ^42,43^. We previously named the resulting His-rich (5.9%) and Tyr-rich (6.5%) sequence as ir8H2, which corresponds to the IDP-sensing domain of Sensor A in Fig. 1B. Fusing ir8H2 to sfGFP and comparing it with r8, we saw a significant improvement in its ability to distinguish mRFP1-(r8)_8_- Ctail condensates from the cytosolic background signal (Fig. 2B). Quantitatively, the average partition coefficient increased by 55% (Fig. 2C; *P*=3.3 ±0.7). Departing from r8 and its variants, we considered ieFLG1, a 200-residue IDP (Fig. S1C) consisting of five repeats of a zwitterionic His-rich (15%) sequence (SYGRHGSDGHGARDSQEHYGQRQHSHGSRDGQYSHSGDRG) designed de novo to match the composition of r8 while augmenting LLPS-relevant features, namely high tyrosine (7.5%) content^7^. ieFLG1 shares little sequence identity to r8 (20.1%) and to ir8H2 (19.8%). When fused to sfGFP, ieFLG1 prominently marked mRFP1-(r8)_8_-Ctail condensates (Fig. 2B), outperforming r8 with a 3.4-fold increase in the average partition coefficient (Fig. 2C; *P*=7.2 ±1.8). These observations demonstrated that the IDP-sensing domain can be evolved to modulate the sensitivity of LLPS-sensors.

We next asked how the fluorescent protein domain of the LLPS-sensor dictates its sensitivity. We set out to test how the surface properties of closely-related green fluorescent proteins (Fig. 3A) influenced sensor performance. Specifically, we focused on surface charge as an accessible variable that is relevant to LLPS dynamics. sfGFP harbors a net charge of -6 and other groups have developed positive and negatively supercharged sfGFP variants ^44^. We selected two of these variants: n20GFP with a net charge of -20, and p15GFP with a net charge of +15. Fusing these sfGFP variants to our IDP-sensing domains, we found that n20GFP abolished the sensitive marking of mRFP1-(r8)_8_-Ctail by ir8H2 (Fig. 3A) and ieFLG1 (Fig. S2), yielding partition coefficients consistently lower than what we measured with sfGFP fusions (*P*=1.8 ±0.4; Fig. 3C and Fig. S3). In contrast, p15GFP greatly improved overall recruitment of ir8H2 into mRFP1-(r8)_8_- Ctail condensates (Fig. 3B), increasing the average partition coefficients tenfold compared with sfGFP (*P*=23.7 ±8.9; Fig. 3C). This enhanced sensitivity was also seen for p15GFP fusions to ieFLG1 (*P*=35.5 ±15.1; Fig. S2-S3) and to a lesser extent for the suboptimal r8 domain (*P*=8.2 ±2.5; Fig. S3). Because the supercharging mutations in p15GFP favored LLPS-relevant Arginine (Arg) residues, we mutated the original eight Arg mutations into lysine (Lys or K). The new sfGFP variant, which we named p15GFPKv, retained the same net charge (Fig. 3A) but lacked an Arg- decorated surface. Interestingly, p15GFPKv still outperformed sfGFP and allowed for sensitive marking of mRFP1-(r8)_8_-Ctail condensates with both ir8H2 (Fig. 3B) and ieFLG1 (Fig. S2), but with a consistent and significant twofold drop in the average partition coefficient compared with p15GFP fusions (*P*=11.3 ±5.0 for ir8H2 and *P*=17.6 ±7.1 for ieFLG1; see quantifications in Fig. S3). These data demonstrated that the sensitivity of LLPS-sensors can be readily tuned by both the net charge and surface chemistry of the fluorescent protein domain. Overall, our quantifications also showed for the first time that the two epidermal LLPS-sensors that we previously deployed in mice, Sensor A and Sensor B, feature optimized fluorescent protein (p15GFP) and IDP-sensing domains —with ir8H2 for Sensor A (Fig. 1B) and ieFLG1 for Sensor B^7^.

**Fig 3.**
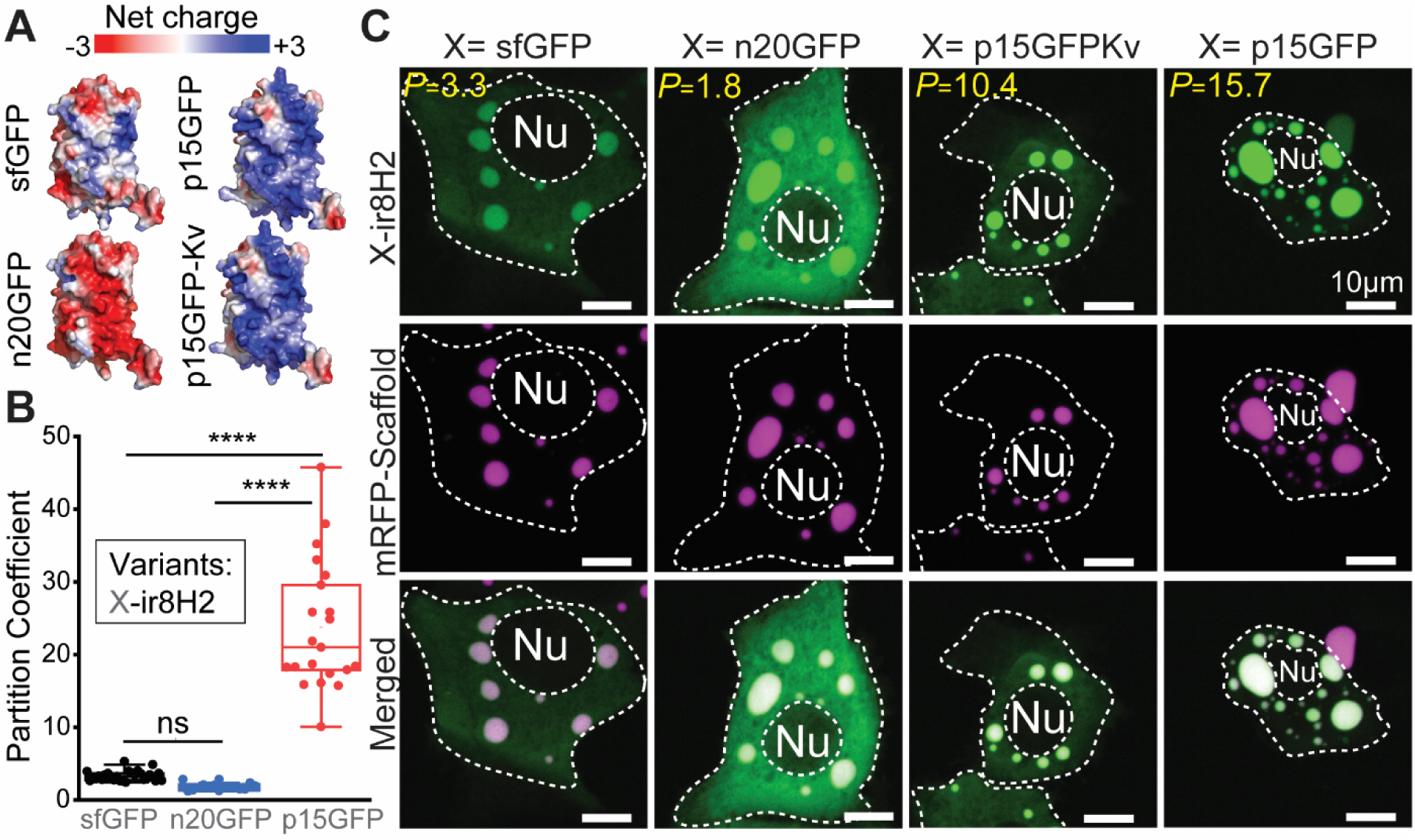
The fluorescent protein domain potently alters the sensitivity of the LLPS-sensing IDP domain. **(A)** AlphaFold2 predictions of the 3D surface of four variants of sfGFP, colored by net charge. **(B-C)** Live-cell images (**B**) and quantifications (**C**) of LLPS-sensor variants partitioning into mRFP1-(r8)_8_-Ctail condensates. The variants share a fixed LLPS-sensing IDP (ir8H2) but differ in the fluorescent protein domain (X). This sensitivity enhancement from p15GFP over all sfGFP variants, including p15GFPKv, was consistent across LLPS-sensing domains (see Fig. S2-S3). n20GFP: sfGFP with a negatively charged surface and -20 net charge. p15GFP: sfGFP with a positively charged surface and a +15 net charge. p15GFPKv: variant of p15GFP wherein surface-exposed Arg residues were mutated to Lys residues. *P*: Partition Coefficient. Nu: nucleus. **(C)** Each data point corresponds to the LLPS-sensor partition coefficient across all sizable condensates in a cell. Asterisks denote statistical significance (p≤0.0001). ns: not statistically significant (p>0.05).

### LLPS-sensors do not drive intracellular phase separation

Despite carrying LLPS-relevant sequence information, optimized LLPS-sensors must remain soluble and diffuse in the absence of the IDP-scaffold or prior to the onset of intracellular LLPS (Fig. 1A). Conceptually, LLPS-sensors should not exhibit concentration-dependent intracellular LLPS. In our extensive in vivo characterization of Sensor A, we previously saw that intracellular Sensor A in the spinous layer, prior to the onset of *FLG* expression and KG assembly in the granular layer, appeared diffuse in the cytoplasm^7^. Moreover, in mice that had undergone *flg* knockdown *in utero,* intracellular Sensor A signal appeared diffuse in all epidermal layers^7^.

To further probe the intracellular behavior of Sensor A in the absence of an IDP-scaffold, we set out to study its intracellular localization as a function of expression levels in HaCATs, contrasting it with mRFP1-(r8)_8_-Ctail as a representative IDP-scaffold. For a rigorous comparison of their concentration-dependent LLPS behavior, we built comparable nuclear concentration reporters for each protein using chromatin (H2B) (Fig. 4A). We sought to avoid concerns related to protein size differences (303 KDa vs 64 KDa) and expected changes in localization (condensates vs diffuse). We used a self-cleavable P2A domain to generate equimolar amounts of these proteins with either a mRFP1-tagged H2B protein (H2BRFP; linked to Sensor A) or a GFP-tagged H2B (H2BGFP; linked to the IDP-scaffold). Using live-cell imaging, we captured the intracellular distribution of each protein and quantified the extent of phase separation in individual cells (Fig. 4B), using the fluorescence intensity of the H2B protein as a proxy for intracellular levels. To account for differences in fluorescent intensity between H2BRFP and H2BGFP, we transformed all H2BRFP measurements to H2BGFP units using our experimentally-determined (3:1) H2BGFP-to-H2BRFP ratio ^7^.

**Fig 4:**
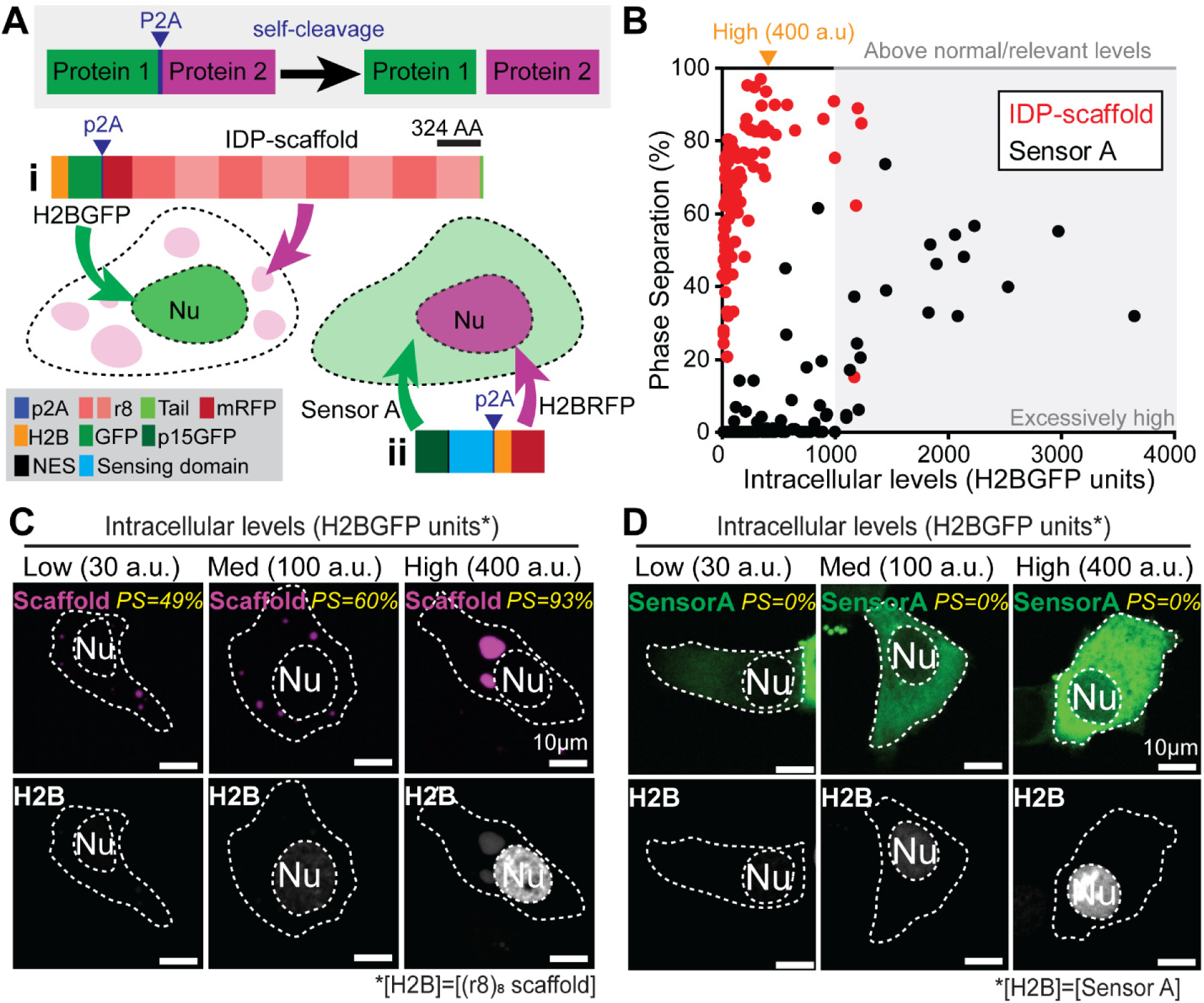
LLPS-sensors do not drive assembly of intracellular condensates. **(A)** Strategy to compare the concentration-dependent phase behavior of a FLG-like IDP-scaffold and its LLPS-sensor (Sensor A) — without co-expression of the scaffold. The P2A self-cleavage mechanism (light gray) enables the use of fluorescent chromatin (H2B) as a consistent storage medium of the intracellular expression levels for both (**i**) mRFP1-(r8)_8_-Ctail and (**ii**) Sensor A. **(B)** Extent of intracellular LLPS for mRFP1-(r8)_8_-Ctail or Sensor A (alone) over a wide range of expression levels. Upon excessive overexpression (gray region), Sensor A aggregated without condensate formation (see Fig. S4). H2BRFP units were converted to H2BGFP units using an experimentally-validated (3:1) H2BGFP-to-H2BRFP ratio ^7^. **(C)** Live-cell images showing condensate formation by mRFP1-(r8)_8_-Ctail (magenta) over the entire range of protein expression levels in (B). **(D)** Live-cell images showing diffuse Sensor A signal (green) across an LLPS-relevant range of expression levels.

mRFP1-(r8)_8_-Ctail exhibited a sharp phase transition at low expression levels (Fig. 4B), with the fraction of IDP-scaffold within liquid-like condensates approaching 100% at high expression levels (∼400 a.u.; Fig. 4C). Condensates were already prominent at low expression levels (∼30 a.u.; Fig. 4C). In contrast, Sensor A did not exhibit concentration-dependent LLPS over a wide concentration range (up to ∼1000 a.u., Fig. 4B). For a handful of cells with abnormally high expression levels (>1000 a.u.), mRFP1-(r8)_8_-Ctail signal within condensates dropped (Fig. 4B) and Sensor A signal clustered into irregular aggregates (Fig. S4) that lacked the liquid-like sphericity of the mRFP1-(r8)_8_-Ctail condensates. We considered this concentration regime as prone to artifacts from transfection. Within the LLPS-relevant range of concentrations, we confirmed that Sensor A at low (30 a.u.), medium, (100 a.u.) and high (400 a.u.) intracellular levels distributed diffusely in the cytoplasm (Fig. 4D). Together with our observations in Fig. 3, these live-cell data demonstrated that optimized LLPS-sensors behave as soluble intracellular proteins that are highly sensitive to the density phase transitions exhibited by target IDP-scaffolds.

### Benchmarking LLPS-sensor performance with ligand-type clients

Biomolecular condensates and their IDP-scaffolds interact with client biomolecules across a broad range of affinities. One prominent example involves ligand-type client proteins that bind one-to-one via a specific domain in the IDP-scaffold^2,4,18,19,45^. We reasoned that fluorescent ligand-type client proteins offered an intriguing alternative to monitor the intracellular LLPS dynamics of target IDP-scaffolds with known binding domains.

We set out to contrast the performance of Sensor A and a fluorescently-tagged client for probing the intracellular LLPS dynamics of FLG-like IDP-scaffolds. We selected a dead variant of the Tobacco Etch Virus protease (dTEVp) fused to sfGFP as our model client (Fig. 5A). We previously reported that sfGFP-dTEVp co-localizes with condensates assembled by mRFP1-cTEV-(r8)_8_- Ctail. This FLG-like IDP-scaffold features the short motif ENLYFQS, which is the known binding and cleavage site (cTEV) for TEVp. As suggested by the droplet-promoting propensity profiles in Fig. 5A, this dTEVp client lacks sequence-level LLPS information. This is in line with our prior observation that sfGFP-dTEVp was excluded from condensates formed by mRFP1-(r8)_8_-Ctail modified with a defective mutant cTEV motif (ENLYFQ<u>R</u>)^7^. While sfGFP-dTEVp is comparable in size to Sensor A, it lacks the disordered conformational dynamics of Sensor A and the target IDP- scaffold (Fig. 5B and Fig. S5).

**Fig 5.**
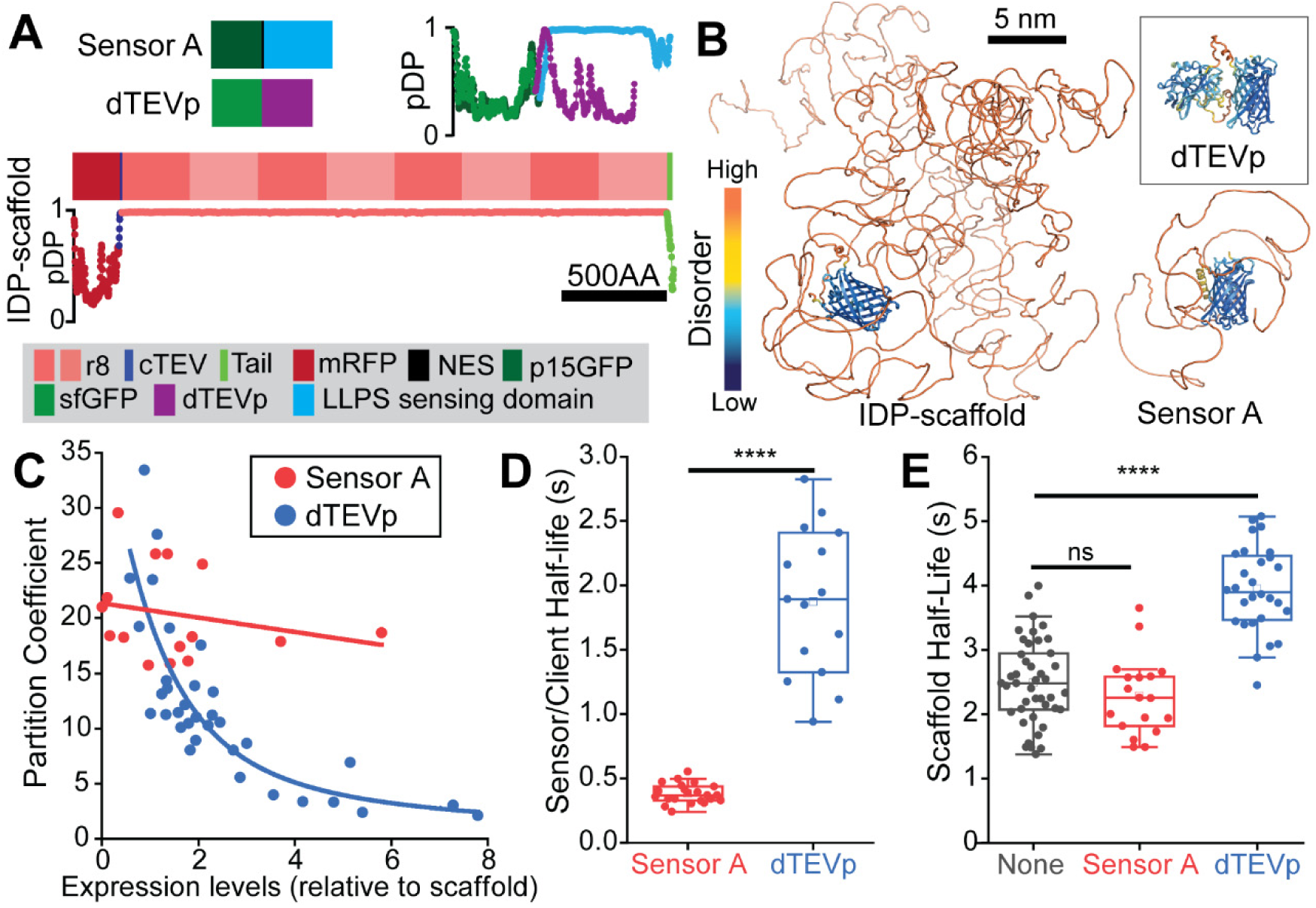
LLPS-sensors faithfully monitor condensate assembly and their liquid-like dynamics. **(A)** Domain architecture and droplet-promoting propensity (pDP; where 1 indicates highest LLPS information content) plots for three constructs: (i) mRFP1-cTEV-(r8)_8_-Ctail, a FLG-like IDP-scaffold engineered to incorporate a cTEV site; (ii) an engineered fluorescent client that binds to cTEV (dTEVp client); and (iii) Sensor A. dTEVp: dead variant of the Tobacco Etch Virus protease. cTEV: cleavage (and binding) site for TEVp. **(B)** AlphaFold2 predictions of the 3D structure and disorder for the three proteins in (A). **(C)** Quantification of Sensor A or dTEVp client partitioning into condensates as a function of their intracellular levels relative to the concentration of mRFP1-cTEV-(r8)_8_-Ctail. Unlike the dTEVp client, Sensor A sensitively marked condensates over a wide concentration range of sensor and IDP-scaffold levels. **(D)** Sensor A and dTEVp client recovery half-lives after internal photobleaching of the corresponding mRFP1-cTEV-(r8)_8_-Ctail condensates. Compared with the scaffold-interacting dTEVp client, Sensor A undergoes relatively unimpeded mixing within condensates. **(E)** IDP-scaffold recovery half-lives after internal photobleaching of the corresponding condensates with and without enrichment of Sensor A or dTEVp as indicated. The client-scaffold interactions slowed down the liquid-like dynamics of the scaffold (i.e., longer scaffold half- life), whereas Sensor A preserved the native liquid-like dynamics of the IDP-scaffold. Raw data in (D)-(E) reanalyzed from supplementary information in ^7^. Asterisks: statistically significant (*=p≤0.05; ****= P≤0.0001). ns: not statistically significant (p>0.05).

Fine control over the intracellular levels of genetically encoded tools remains challenging, suggesting a requirement for intracellular LLPS-sensors to perform well over a wide concentration range. Here, we asked how the relative abundance of sensor/client to IDP-scaffold influenced the sensitive marking of intracellular condensates. We took advantage of the variable expression levels of mRFP1-cTEV-(r8)_8_-Ctail and either Sensor A or sfGFP-dTEVp upon transfection in HaCATs. At low intracellular levels, both Sensor A and sfGFP-dTEVP showed sensitive marking of mRFP1-cTEV-(r8)_8_-Ctail condensates with comparably high partition coefficients (*P*∼20-30; Fig. 5C). Sensor A sustained its high sensitivity even when expressed at higher levels than the IDP-scaffold (Fig. 5C). In contrast, the partition coefficient of sfGFP-dTEVp decreased precipitously as the client-to-scaffold levels exceeded a ratio of 1 (Fig. 5C), pointing to the concentration-dependent saturation of dTEVp-bound cTEV sites within condensates. We will consider the impact of the number of cTEV sites per IDP-scaffold later in this paper. These data suggested that ligand-type clients may have subpar performance in monitoring early intracellular LLPS dynamics dictated by the gain of IDP-scaffold expression —like *FLG* upregulation in the granular layer of human skin.

At late stages of intracellular LLPS, we previously reported that the liquid-like dynamics of mRFP1-cTEV-(r8)_8_-Ctail within condensates is sensitive to the binding of sfGFP-dTEVp but not Sensor A ^7^. We revisited these supplemental data and reanalyzed it to directly contrast the behavior and impact of Sensor A and sfGFP-dTEVp (Fig. 5D-E). To do this, we considered internal fluorescence recovery after photobleaching (FRAP) measurements, this time as unnormalized recovery half-life values. These internal photobleaching experiments allowed us to quantify the relative diffusion dynamics of the IDP-scaffold and our fluorescently-tagged tools within condensates. Sensor A showed greatly accelerated diffusion compared with sfGFP-dTEVp, exhibiting full recovery in less than a second (Fig. 5D). sfGFP-dTEVp exhibited an average recovery half-life that was four-fold longer than that of Sensor A (Fig. 5D), with values comparable to the half-life of the IDP-scaffold (Fig. 5E) despite the large size difference between client and scaffold (Fig. 5B). These divergent dynamics reinforced the notion that Sensor A interacts very weakly with the IDP-scaffold, even within condensates. The surprisingly slow diffusion dynamics of dTEVp suggested substantial binding to the IDP-scaffold, which translated into a significant slowing down of IDP-scaffold dynamics (Fig. 5E). On the other hand, prominent Sensor A recruitment to condensates did not significantly alter the IDP-scaffold half-life values, which were indistinguishable from control condensates (Fig. 5E). Overall, our analyses showed that the ultraweak interactions between LLPS-sensor and IDP-scaffold allow for sensitive recruitment in a concentration-independent manner and without negative impact to IDP-scaffold liquid-like dynamics within condensates.

### LLPS-sensors innocuously probe the onset of intracellular LLPS

Our initial observations focused on late-stage intracellular LLPS, which feature prominent biomolecular condensates that are amenable for FRAP assays. However, epidermal LLPS- sensors are expressed in vivo prior to the onset of LLPS. We turned our attention towards testing how Sensor A and sfGFP-dTEVp influenced the concentration-dependent onset of intracellular LLPS by a range of FLG-like IDP-scaffolds. To avoid confounding effects from differing intracellular ratios of IDP-scaffold and sensor/client, we deployed our P2A system to produce each IDP-scaffold and sensor/client at equimolar levels (Fig. 6A).

**Fig 6.**
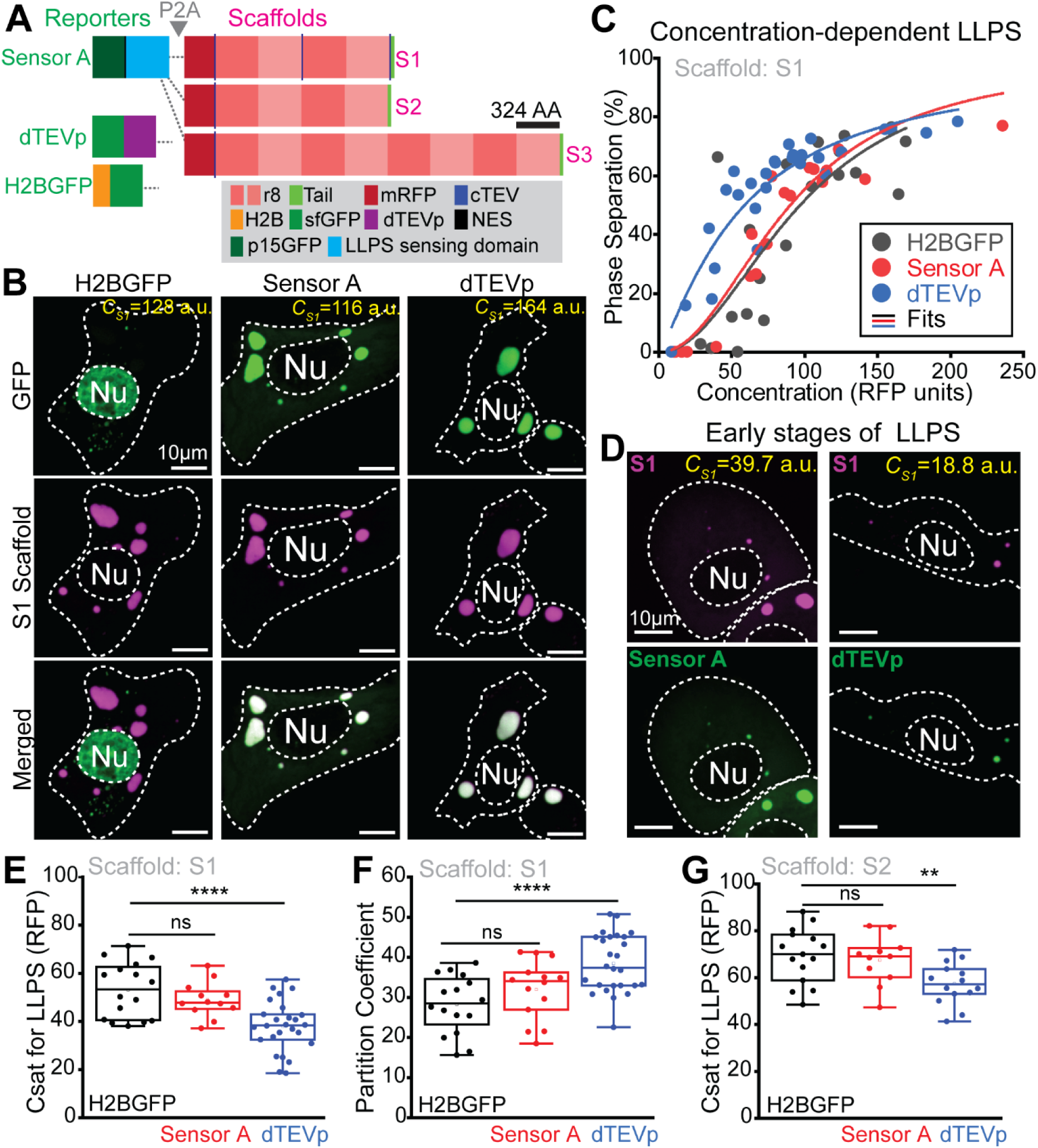
LLPS-sensors innocuously probe the intracellular phase separation dynamics of IDP-scaffolds. **(A)** Strategy to test the impact of Sensor A co-expression on the intracellular LLPS dynamics driven by three IDP-scaffolds (S1-S3). Self-cleavable p2A linkage ensures that Sensor A is abundantly present as the IDP-scaffold accumulates beyond its saturation concentration (C_sat_ for LLPS). This strategy was extended to a fluorescent ligand-type client (dTEVp) and a non-interacting control protein (nuclear H2BGFP), all p2A-linked to S1-S3. **(B)** Live-cell images H2BGFP, Sensor A and dTEVp client localization in cells with well-established S1 condensates and similar levels of S1 scaffold (∼120-160 RFP units). **(C)** Phase separation of the S1 scaffold as a function of its intracellular levels (RFP units) in cells with equivalent expression levels of H2BGFP (control), dTEVp client or Sensor A. **(D)** Live-cell images of Sensor A or dTEVp recruitment to condensates in early LLPS. **(E)** Critical concentrations for LLPS of S1 scaffold the presence of H2BGFP (black), Sensor A (red), and dTEVp (blue). **(F)** Partition coefficients of S1 scaffold in the presence of H2BGFP (black), Sensor A (red), and dTEVp (blue). **(G)** Critical concentrations for LLPS of S2 scaffold in the presence of H2BGFP (black), Sensor A (red), and dTEVp (blue). Asterisks denote statistical significance (** = P≤0.01; **** = P≤0.0001). ns: not statistically significant (p>0.05).

Our IDP-scaffolds in Fig. 6A varied in length (4 vs 8 repeats of r8) and the number of cTEV sites (1 vs 3). Using our live-cell imaging approach in HaCATs, we quantified the IDP-scaffold concentration outside (dilute phase) and within condensates for three IDP-scaffolds (S1-S3 in Fig. 6A) in the presence of Sensor A, sfGFP-dTEVp or the nuclear-localized H2BGFP as a non-interacting control protein (Fig. 6A). We treated the dilute phase concentration as a proxy for the C_sat_. Comparing the S1 and S2 scaffolds in the H2BGFP controls, we were surprised to notice that the two additional instances of the short cTEV motif in S1 (<1% of its residues) led to a significant drop in the C_sat_ and increased its partition coefficient (Fig. S6A-B). Interestingly, when co-expressing with sfGFP-dTEVp, S1 condensates showed a nearly four-fold increase in sfGFP- dTEVp recruitment compared with S2 condensates (Fig. S6C). Sensor A partitioning was insensitive to the number of cTEV sites in these IDP-scaffolds, showing similar partition coefficients into condensates formed by S1 and S2 (Fig. S6C). These data suggested that even large IDP-scaffolds may be sensitive to small sequence modifications required for engineered ligand-type clients.

Next, we examined whether Sensor A and sfGFP-dTEVp shifted the LLPS-relevant metrics of the S1 scaffold. Live-cell imaging showed robust Sensor A and dTEVp recruitment to large S1 condensates, while our control H2BGFP protein localized to the nucleus as expected (Fig. 6B). Quantifying the concentration-dependent intracellular LLPS of S1, we saw that co-expression with dTEVp displayed a leftward shift in the extent of LLPS as a function of S1 levels when compared with H2BGFP (Fig. 6C). In this system, the equimolar synthesis of S1 and dTEVp prevented saturation of the three cTEV sites in S1, resulting in sensitive marking of nascent condensates (Fig. 6D). In line with the observed concentration-dependent shift, the C_sat_ for S1 in the presence of dTEVp dropped significantly (Fig. 6E) while the scaffold partition coefficient increased (Fig. 6F) compared with H2BGFP controls. In notable contrast, Sensor A did not disrupt any of these key intracellular LLPS metrics for S1 (Fig 6C, E-F).

We repeated these analyses with the S2 scaffold, which is identical to S1 except that it features only one cTEV motif. We were motivated to test if limiting the client-scaffold interaction to one short motif rescues the client-linked disruptions of the endogenous LLPS dynamics. Sensor A and dTEVp sensitively marked S2 condensates (Fig. S7A). We confirmed that Sensor A did not shift the concentration-dependent LLPS of S2 (Fig. S7B), the C_sat_ (Fig. 6G), and the partition coefficient (Fig. S7C). dTEVp still noticeably shifted the concentration-dependent LLPS of S2 (Fig. S7B), significantly reducing the C_sat_ (Fig. 6G) and increasing the expected partition coefficient (Fig. S7C). We then repeated these experiments with S3, which is similar to S2 but features double the number of r8 repeats (Fig. 6A). For this large IDP-scaffold, Sensor A and dTEVp performed similarly, both faithfully reporting the LLPS dynamics seen for the H2BGFP controls (Fig. S8). We note, however, that our prior observations on IDP-scaffold diffusion dynamics within condensates involved this S3 scaffold (Fig. 5E), suggesting that even large IDP-scaffolds may remain susceptible for disruption within the protein-rich condensate environment. Overall, these data demonstrated that LLPS-sensors consistently enabled high-fidelity intracellular probing of diverse IDP-scaffolds and that ligand-type clients may be suitable probes for a subset of IDP-scaffolds.

### Ultraweak interactions for high-fidelity probing of intracellular LLPS

Our findings for Sensor A and our dTEVp client suggest that the interaction mode with the IDP- scaffold is a key variable in the evolution of tools for high-fidelity intracellular probing of the LLPS dynamics of IDP-scaffolds and their condensates (Fig. 7). Classical ligand-type clients bind to specific motifs along IDP-scaffolds with moderate-to-high affinity. Alternatively, LLPS-sensors engage in the same ultraweak intermolecular interactions that govern the LLPS of an IDP-scaffold (Fig. 7A). This unique mode of interaction enables the sensor to engage weakly with the regions of the IDP-scaffold that encode LLPS-relevant information, with the bulk of transient scaffold- sensor interactions occurring within condensates. We demonstrate that these differences in interaction modes have critical implications for probing the concentration-dependent LLPS of the IDP-scaffold (Fig. 7B).

**Fig 7:**
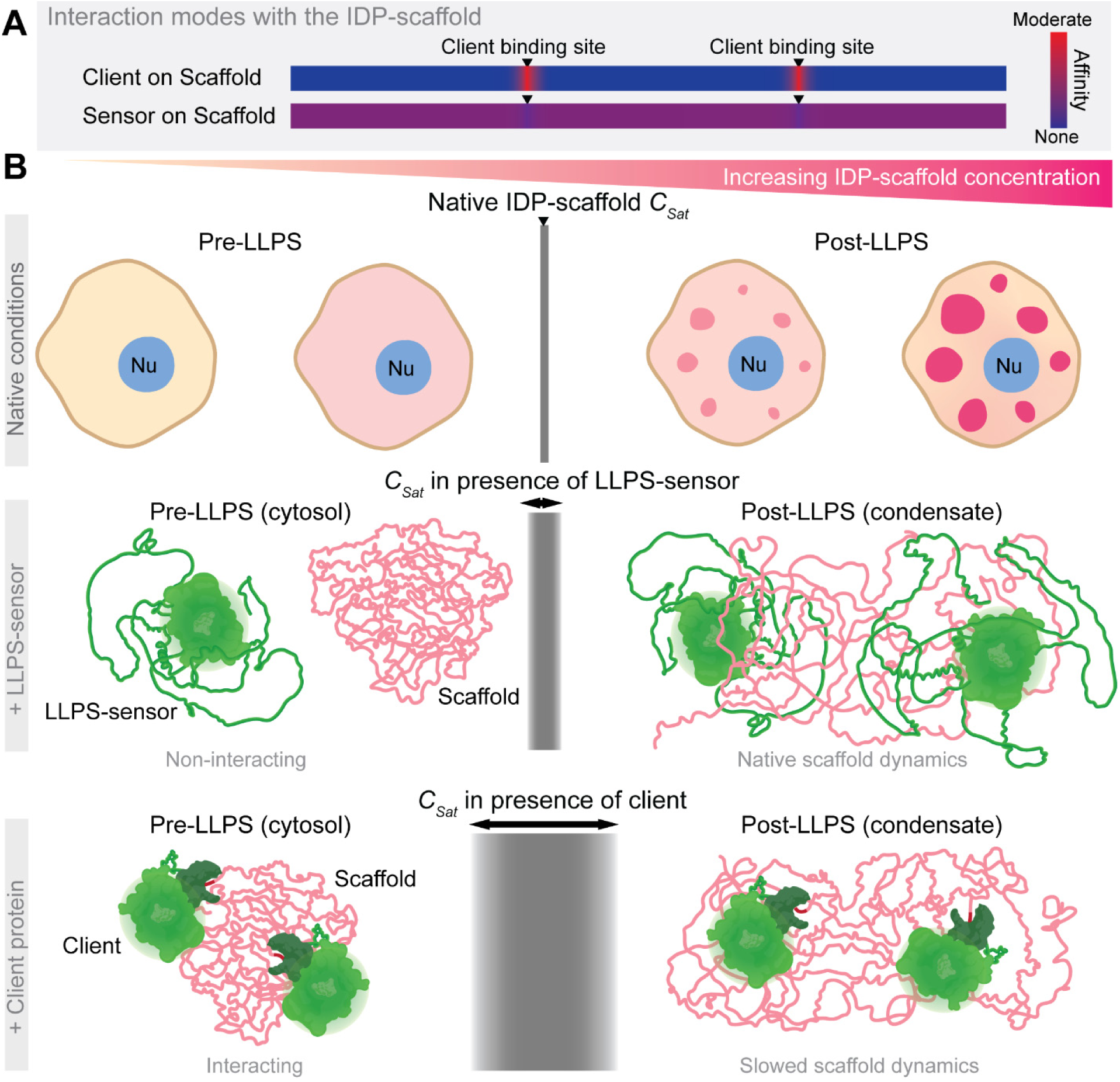
Ultraweak and phase-activated sensor-scaffold interactions enable sensitive and innocuous probing of intracellular condensate assembly and LLPS dynamics. **(A)** Ligand-type clients and LLPS-sensors differ in their interaction modes with IDP-scaffolds, distinguished by the affinity level and locations of the protein-protein interactions. **(B)** The resulting interaction dynamics help explain the unique ability of LLPS-sensors to faithfully monitor and report the native (top panel) LLPS dynamics of IDP-scaffolds. Prior to the saturation concentration (C_sat_) of the IDP-scaffold, LLPS-sensors (middle panel) do not interact with IDP-scaffolds that exist in a partially collapsed conformation reflective of favored intermolecular interactions ^46^, avoiding shifts to the native C_sat_ of the IDP-scaffold. Upon phase separation above C_sat_, IDP-scaffolds open to intermolecular interactions and weakly but multivalently engage with LLPS-sensors to sustain their LLPS-sensitive enrichment within nascent and established condensates —irrespective of the relative sensor levels. These phase-activated sensor-scaffold interactions are weak and highly dynamic, preventing shifts in the scaffold liquid-like dynamics within condensates. Unlike LLPS-sensors, co-expressed ligand-type clients (bottom panel) may bind to the IDP-scaffold with moderate affinity (i.e., even low μM level), allowing for interaction in the cytosol before the scaffold accumulates and phase separates above the native C_sat_. These substantial interactions can cause major shifts in the IDP-scaffold C_sat_, inhibiting or inducing (as we saw for the dTEVp client in Fig. 6) intracellular phase separation, and inevitably slow down scaffold dynamics within assembled condensates.

Moderate-to-high affinity interactions between ligand-type clients and IDP-scaffolds may not discriminate the biophysical features of the IDP-scaffold before and after intracellular LLPS (Fig. 7B). Rather, ligand-type clients that bind to the IDP-scaffold with high (1-100 nM) or moderate (>0.1-10 uM) affinity are likely to substantially interact with the IDP-scaffold in the dilute phase (Fig. 7B). Our data demonstrate that these interactions can shift the underlying intracellular LLPS dynamics of select IDP-scaffolds. For IDP-scaffolds that tolerate weak binding in the dilute phase, like S3 in Fig. S8, we showed that the liquid-like dynamics within condensates remain particularly sensitive to this binding (Fig. 5E and Fig. 7B). In the case of our sfGFP-dTEVp client, its interactions with a range of IDP-scaffolds favored the assembly of dense condensates. Generally, we surmise that ligand-type clients may alter the solubility and conformational dynamics of IDP- scaffold chains to either increase or decrease intracellular LLPS propensity ^45,47^.

LLPS-sensor interaction with the target IDP-scaffold is conditional upon the enrichment of IDP- scaffold within condensates, where they reside at high density and in an extended conformation (Fig. 7B). Below the C_sat_ of the IDP-scaffold, the IDP chains are diffuse in the intracellular space at low concentrations that are unlikely to allow for substantial interaction with the LLPS-sensor — considering affinities in the 100 μM range for self-interacting IDPs. In this dilute regime, LLPS-prone IDP-scaffolds exist in a collapsed state dominated by intramolecular scaffold interactions, reducing the likelihood of heterotypic scaffold-sensor interactions^46,48^. Without substantial interactions in the dilute phase, the biophysical and conformational properties of the IDP-scaffold are not perturbed prior to the onset of phase separation. Upon a phase transition, however, IDP- scaffold chains accumulate at high density within condensates —often well above 100 μM ^7^— and adopt an extended conformation^46,48^, promoting multivalent sensor-IDP-scaffold interactions that drive sensor accumulation within condensates. The interactions in the dense phase remain highly dynamic, preventing LLPS-sensors from interfering with the internal liquid-like dynamics of condensates.

Beyond our proof-of-principle work with epidermal LLPS-sensors, we acknowledge the challenge of engineering LLPS-sensors for target IDP-scaffolds and their condensates. The evolution of engineered IDPs as bona fide LLPS-sensors demands a difficult balance between maximizing sensitive marking of target condensates while minimizing strong heterotypic interactions, which shift the LLPS dynamics of IDP-scaffolds in multicomponent condensates ^48,49^. Encouragingly, the engineering of IDP-sensing domains may accelerate with rapid progress in the sequence-level prediction of LLPS information in IDPs ^8,50^ and in the computational prediction of their conformational and LLPS dynamics ^51–53^. These protein engineering efforts are poised to benefit from a growing understanding of the LLPS grammar in IDP-scaffolds ^37–39,42,43,54^. The orthogonality between IDP-driven biomolecular condensates remains poorly understood, which raises questions about the ability of optimized LLPS-sensors to distinguish between competing condensates. In keratinocytes, Sensor A without a nuclear export signal partitioned weakly into nucleoli, suggesting preferential marking of FLG condensates ^7^. However, LLPS-sensors fueled by ultraweak multivalent interactions may lack molecular information to distinguish epidermal condensates assembled by closely-related FLG paralogs, such as the RPTN granules that appear to coexist with FLG condensates^6^. In specific intracellular contexts, this crosstalk may facilitate comparative analyses of related condensates with a shared LLPS-sensor.

### Outlook

We demonstrated that genetically encoded LLPS-sensors can enable the high-fidelity biophysical probing of IDP-driven biomolecular condensates. Complementing our studies of epidermal LLPS- sensors in mice, our new data convincingly show the potential to evolve biocompatible sensors with four key features: (1) highly tunable sensitivity, (2) concentration-independent partitioning into target biomolecular condensates, (3) faithful monitoring of the underlying concentration-dependent LLPS dynamics, and (4) innocuous probing of the liquid-like dynamics within condensates. While we focused on tools for biophysical probing, we note that LLPS-sensors can feature additional functional domains, such as enzymes that enable biomolecular characterization of condensates via proximity proteomics ^55^. LLPS-sensors fused to other proteins of interest may functionalize and drug endogenous condensates for therapeutic outcomes. In our efforts to evaluate sensor performance, we exposed the challenge of engineering ligand-type clients for probing the intracellular LLPS of IDP-scaffolds with high fidelity. We expect that the simplicity and generalizability of this approach will fuel interest in evolving novel ligand-type clients towards achieving minimal disruption of the endogenous LLPS dynamics. Given the enigmatic involvement of diverse IDP-scaffolds in physiological and disease mechanisms, we propose that an array of advanced biomolecular tools will be needed to rigorously and innocuously probe native and engineered condensates. We hope that our findings and conceptual progress will stimulate innovations that revolutionize the live-cell probing of IDP-scaffolds and their assemblies.

## Materials and Methods

### Sequence-level prediction of disorder and LLPS-relevant regions

To profile areas of intrinsic disorder in FLG, FLG-like scaffolds and engineered sensor constructs, we used DISOPRED3 ^56^ from the University College London PSIPRED Workbench (http://bioinf.cs.ucl.ac.uk/psipred/). Scores above 0.5 were considered predictive of intrinsic disorder. To expose LLPS-relevant regions (i.e., those rich in features involved in LLPS), we calculated droplet-promoting scores using FuzDrop ^8^(https://fuzdrop.bio.unipd.it/predictor). Generally, regions with scores at or above 0.60 are assumed to be part of droplet-promoting regions ^8^. In our interpretation of this tool, which tends to overestimate the LLPS propensity of IDPs, we considered scores above 0.6 as segments rich in LLPS-relevant sequence features. When comparing the percent sequence identity of r8 and IDP-sensing domains, we aligned the relevant sequences using Clustal Omega ^57^. To gage conformational dynamics, we generated AlphaFold2 predictions using ColabFold and colored the resulting structures using pLDDT scores as proxy for intrinsic disorder ^58,59^.

### Synthesis of repetitive DNAs encoding FLG-like IDP scaffolds

To assemble FLG-like (r8)_n_ IDP-scaffolds, we used recursive directional ligation by plasmid- reconstruction (Pre-RDL) with a modified pET-24a(+) vector featuring a Gly-stop-stop-stop sequence as previously described ^7^. We previously reported successful iterative rounds of Pre- RDL to build genes with two, four or 8 repeats of r8 —with and without the C-terminal tail domain of human FLG. For IDP-scaffolds designed to interact with dTEVp (as in Fig. 6), we used Pre- RDL to add the short cTEV domain to the C-terminus of (r8)_2_, resulting in (r8)_4_ constructs with two dTEVp-binding sites: a C-Terminal cTEV site and one internal cTEV linking two (r8)_2_ repeats. For mammalian expression, we subcloned all fully assembled repeat genes into published pMAX vectors (Amaxa) encoding mRFP1, mRFP1-cTEV, H2BGFP-P2A-mRFP1 or H2BGFP-P2A- mRFP1-cTEV ^7^. Fusion of our cTEV-containing (r8)_n_ genes to constructs with mRFP1-cTEV resulted in IDP-scaffolds with either one or three cTEV sites. To build genes encoding IDP- scaffolds with equimolar co-expression of LLPS-sensor or client (related to Fig. 6A), we modified our published pMAX vectors encoding H2BGFP-P2A-mRFP1-tagged FLG-like IDP-scaffolds to replace the H2BGFP domain with sequence-verified genes encoding Sensor A or a sfGFP-dTEVp client. See Table S4 for protein sequences of all IDP-scaffold constructs. We previously validated the equimolar synthesis of individual P2A-linked proteins by examining H2BRFP-(P2A)-H2BGFP and H2BGFP-(P2A)-H2BRFP constructs ^7^.

### Synthesis of LLPS-sensor variants and dTEVp-based client

We previously reported the synthesis of mammalian-optimized genes encoding three candidate IDP-sensing domains (r8, ir8H2, and ieFLG1; see Table S1 for sequence details), and four candidate fluorescent protein tags: sfGFP and three supercharged variants of (n20GFP, p15GFP, and p15GFPkv) —all featuring a C-terminal short nuclear export signal (LELLEDLTL) ^36^. Using compatible restriction sites in our library of pMAX vectors, we assembled genes that fused each candidate IDP-sensing domain to each fluorescent protein. The pMAX vector encoding the sfGFP-tagged dTEVp client (see Table S3 for sequence details) was previously reported ^7^. For constructs incorporating P2A domains, we PCR-amplified Sensor A or sfGFP-dTEVp adding compatible restriction sites to clone them into our published H2BGFP-(P2A)-H2BRFP vector, replacing the parent H2BGFP domain. These sequence-verified constructs were used directly (Fig. 4A-B) or as cloned genes for the synthesis of constructs with P2A-linking of Sensor/Client and IDP-scaffolds as described above (related to Fig. 6A).

### Quantitative assessment of LLPS-sensor performance

To analyze the intracellular behavior of FLG-like IDP-scaffolds and related tools, we transfected genes of interest into HaCATs and performed live-cell imaging as previously described ^7^. Briefly, using spinning disk confocal microscopy, we generated max intensity projections to quantify key LLPS-relevant metrics. Specifically, using ImageJ we measured the extent of intracellular LLPS by an IDP-scaffold as the percentage of total (background-corrected) fluorescent signal residing within spherical condensates. Similarly, we quantified the partition coefficient of both IDP- scaffolds and our tools (LLPS-sensor designs and a ligand-type client) as the ratio of the (background-corrected) fluorescent signal intensity within condensates to the signal in the cytosol. For cells with condensates, the background-corrected cytosolic signal of IDP-scaffolds served as a proxy for the saturation concentration (C_sat_). When possible, we estimated intracellular concentration using H2B reporter signal adjusted by total cell area to sensitively and consistently measure concentration across divergent proteins of interest. Whenever we observed a concentration-dependent increase in the extent of LLPS (e.g., Fig. 6C), we applied a logistic fit [y = (−100/(1 + (x/x0)^P)) + 100], as expected for a phase transition^7^ using OriginPro.

### Statistical analyses

Statistical significance indicates that we rejected, with a confidence greater than 0.05 (i.e., p < 0.05), the null hypothesis that the difference in the mean values between two datasets equaled zero. To perform this hypothesis testing, we ran two-sample t tests using OriginPro. In all cases, we verified that the statistical differences did not depend on the assumption of equal variance (Welch-correction) between samples.

## Acknowledgments

FGQ holds a Career Award at the Scientific Interface from Burroughs Wellcome Fund. FGQ acknowledges support from the National Institutes of Health (NIH) through DP2 award 1DP2GM149749. ARCA acknowledges support from the National Institute of Arthritis and Musculoskeletal and Skin Diseases of the NIH under award number F31AR081697.

## Supplemental figures

**Fig S1:**
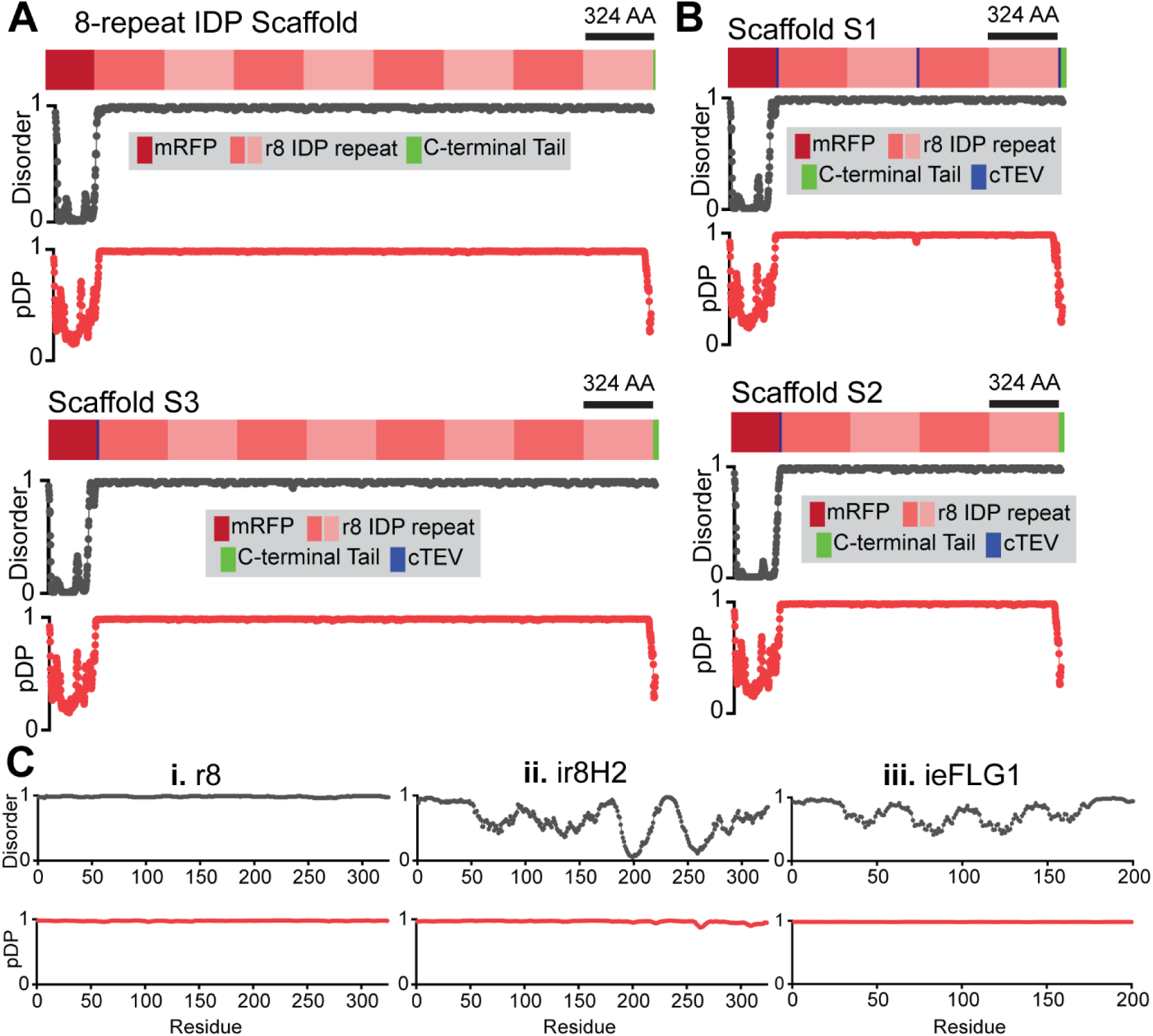
Architecture and features of FLG-like IDP-scaffolds and related LLPS-sensing IDP domains. **(A-B)** Domain architecture and corresponding disorder (where 1=disorder) and droplet-promoting propensity (pDP; where 1 indicates highest LLPS information content) plots for model IDP-scaffolds with **(A)** eight repeats of the r8 domain of human FLG, **(B)** four (S1 and S2) to eight (S3) repeats of r8 and one (S2-S3) to three (S1) cTEV sites. See Fig. 1C for FLG architecture. Note that r8 is ∼95% identical to other repeat domains in FLG (e.g., r1-r7; r9-r10), hence the selection of a specific FLG repeat is inconsequential for the engineering of FLG-like IDP-scaffolds. **(C)** Disorder and pDP plots per residue for (i) r8 and related LLPS-sensing IDP domains (ii) ir8H2 and (iii) ieFLG1. Disorder prediction calculated with *DISOPRED3* (Supplementary Ref. 1) and pDP calculated with FuzDrop (Supplementary Ref. 2).

**Fig S2:**
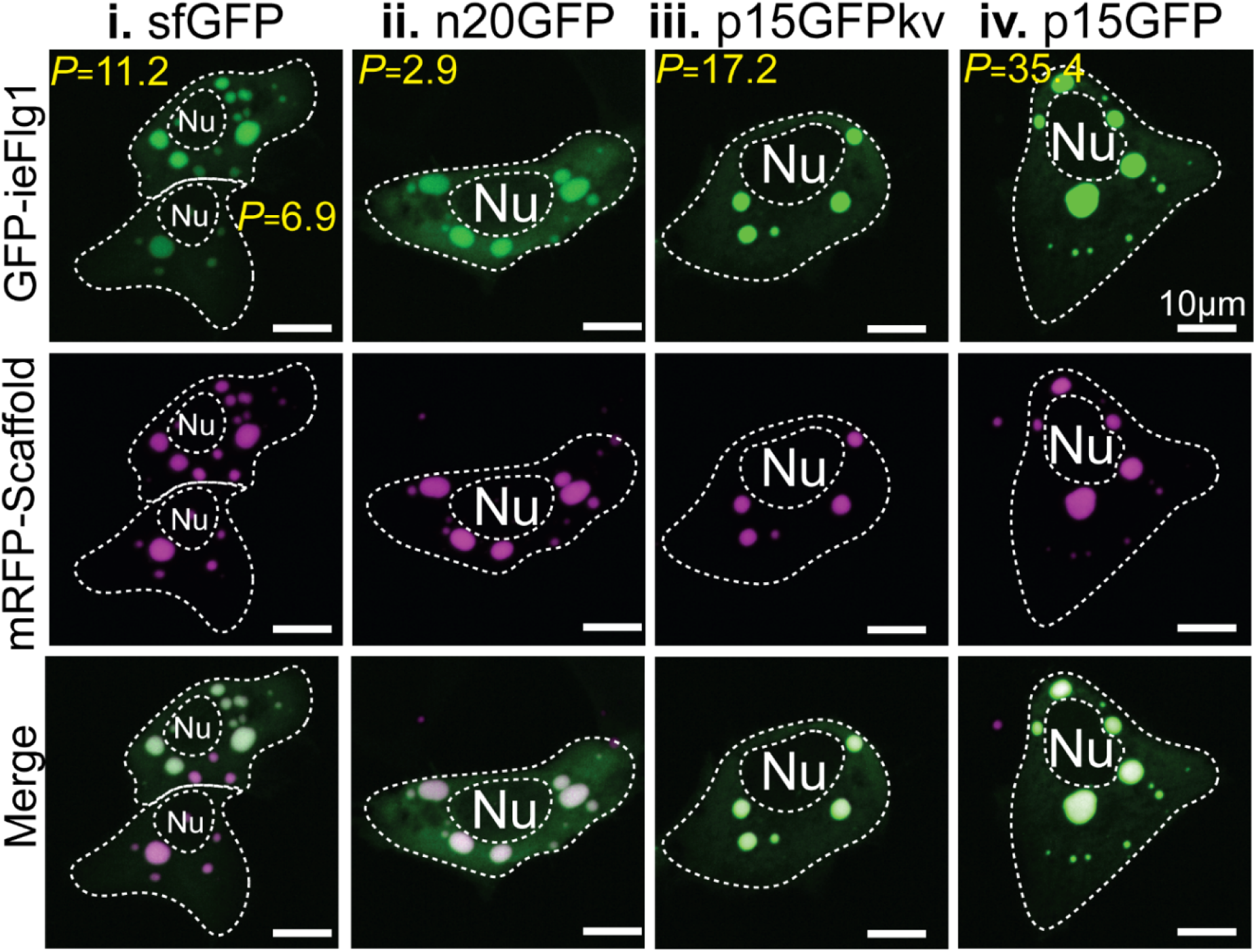
Representative live-cell images comparing the intracellular partitioning of ieFlg1-bearing LLPS-sensors based on the type of GFP-variant. Related to quantifications in Fig. S3 and in line with the observations for ir8H2-bearing LLPS-sensors in Fig. 3B-C and Fig. S3. mRFP-Scaffold: 8-repeat IDP scaffold as shown in Fig. S1A. *P* indicates the partition coefficient for the corresponding FP-ieFLG1 constructs in those specific cells.

**Fig S3:**
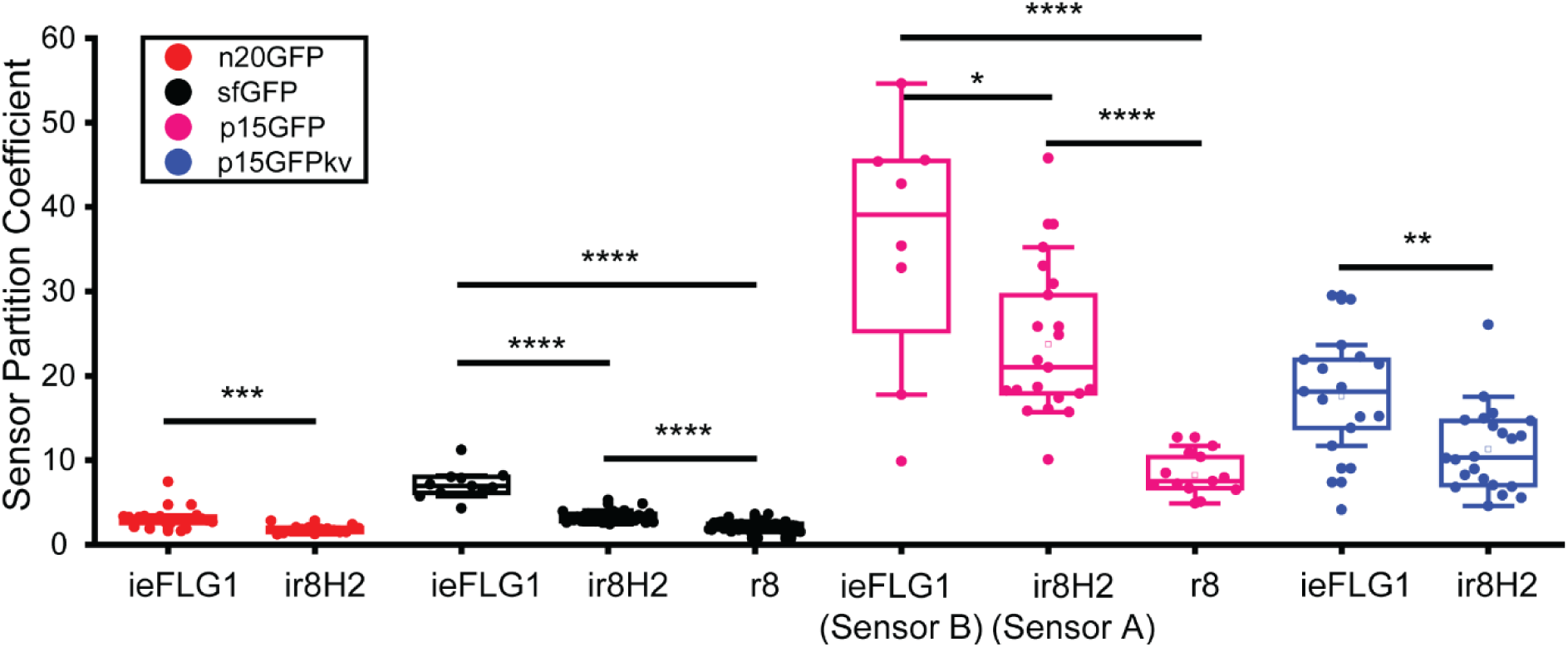
Tunable intracellular partition coefficient across LLPS-sensor designs. Quantitative analysis of the partition coefficient for all tested LLPS-sensor variants, varying IDP-sensor and fluorescent protein domains. These data correspond to sensor partitioning into intracellular condensates formed by the 8-repeat IDP-scaffold of Fig. S1A. These quantifications complement the images and data from Fig. 2, Fig. 3 and Fig. S2. Asterisks denote statistical significance (* = P≤0.05; ** = P≤0.01; *** = P≤0.001; **** = P≤0.0001).

**Fig S4:**
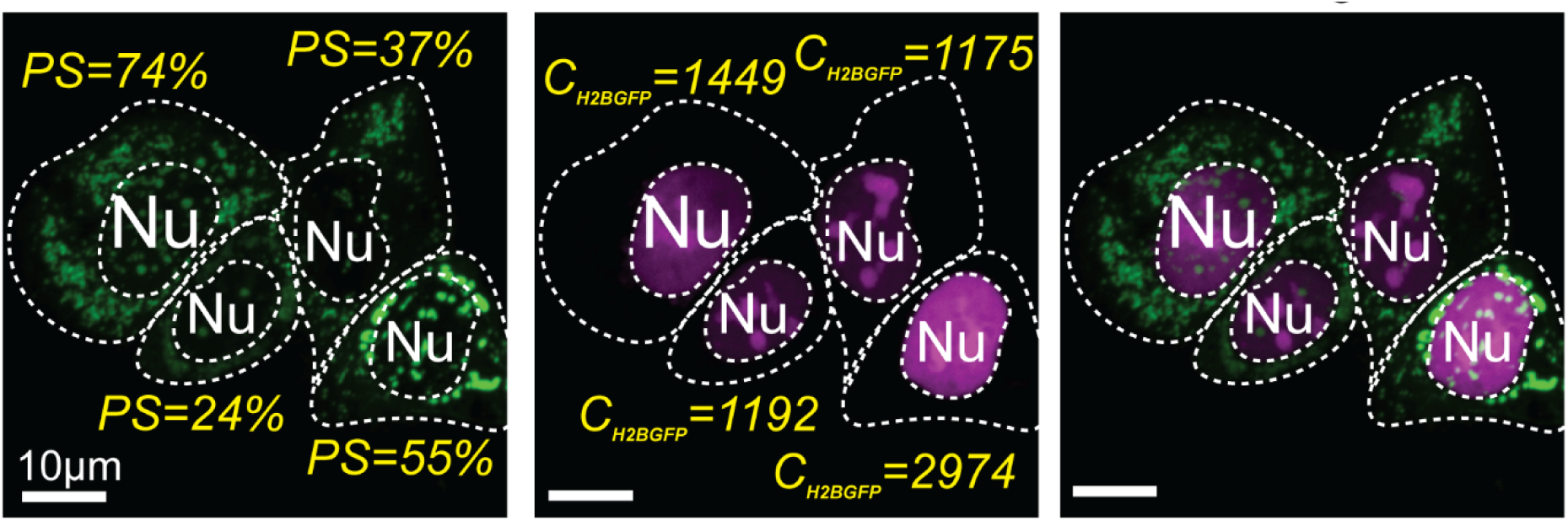
Excessively high intracellular levels of Sensor A results in aggregation rather than phase separation. Live-cell images with corresponding concentration (*C; H2BGFP units*) and percent “phase separation” (PS) measurements for cells expressing abnormally high levels of Sensor A. These levels are not usually achieved during our typical transfection conditions (e.g., when combining scaffold and sensor) and far exceed what we have previously reported in mice upon *in utero* transduction of developing keratinocytes with lentivirus (Supplementary Ref 3). Under these conditions, Sensor A signal revealed irregular clumps that differed morphologically from the liquid-like spherical condensates marked by Sensor A in the presence of (r8)_n_ IDP-scaffolds (for n≥4), FLG (in primary human keratinocytes) and flg (in vivo, in mouse skin). While distinct from phase separation, we chose to quantify and report PS to capture the segregation of fluorescent signal that only occurs when Sensor A is expressed at these abnormally high levels. *C* is reported as *C_H2BGFP_*, converting *C_H2BRFP_* to *C_H2BGFP_* to account for our experimentally-determined 1:3 H2BGFP-to-H2BRFP ratio (Supplementary Ref 3).

**Fig S5:**
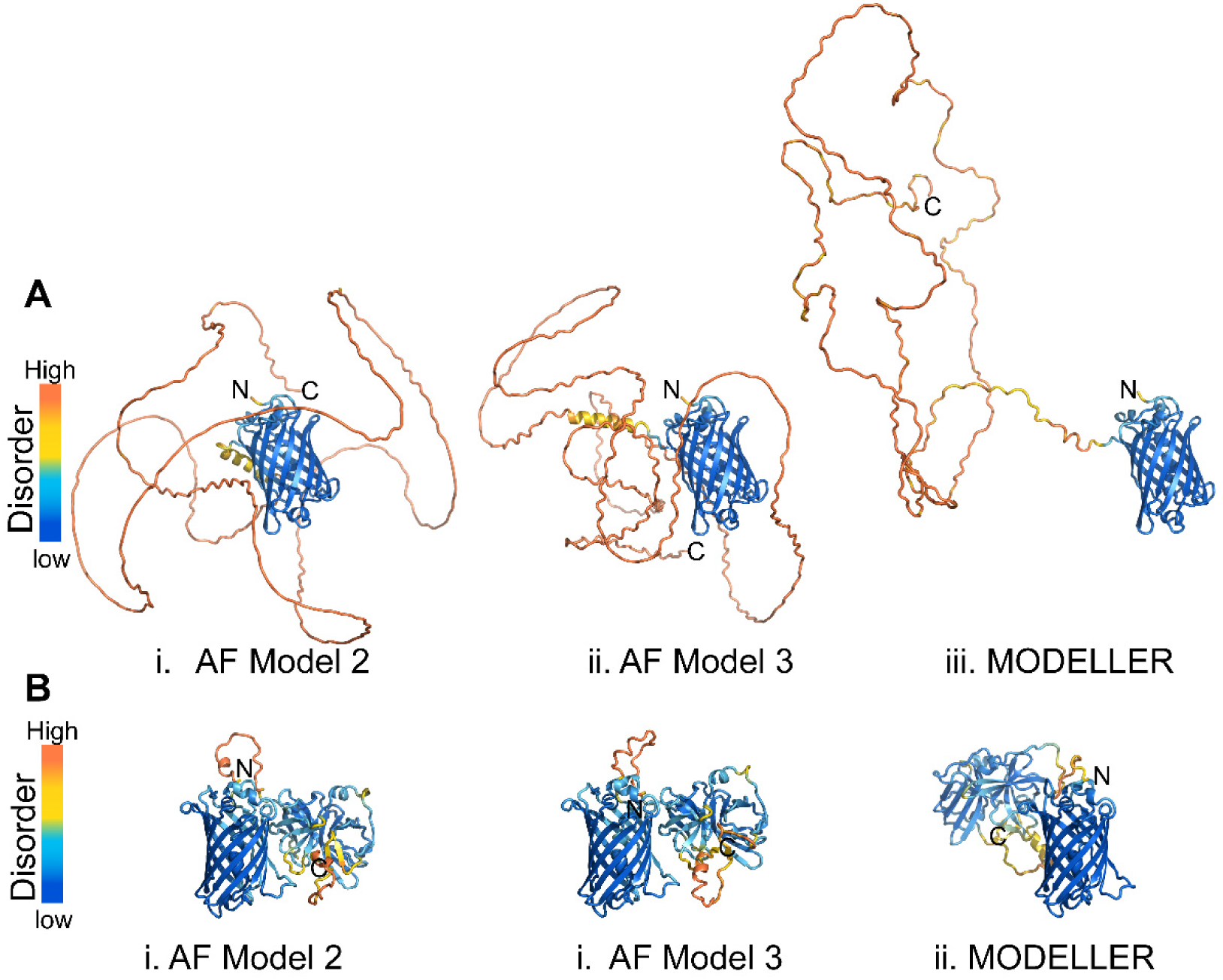
Distinct conformational dynamics between Sensor A and a ligand-type client. **(A-B)** Three snapshots **(i-iii)** of predicted 3D structures for **(A)** Sensor A and **(B)** our sfGFP-dTEVp client. **(A)** The IDP-sensor domain in Sensor A is predicted to sample widely divergent disordered conformations. In contrast, **(B)** the conformational dynamics of the two-domain client protein are largely restricted to small fluctuations in the disordered linker segment. Models (i)-(ii) were predicted by AlphaFold2 from complete protein sequences. Model (iii) was predicted through homology modeling with Modeller (Supplementary Ref 4) from the separate predictions for each of the two domains in either the sensor or client.

**Fig S6:**
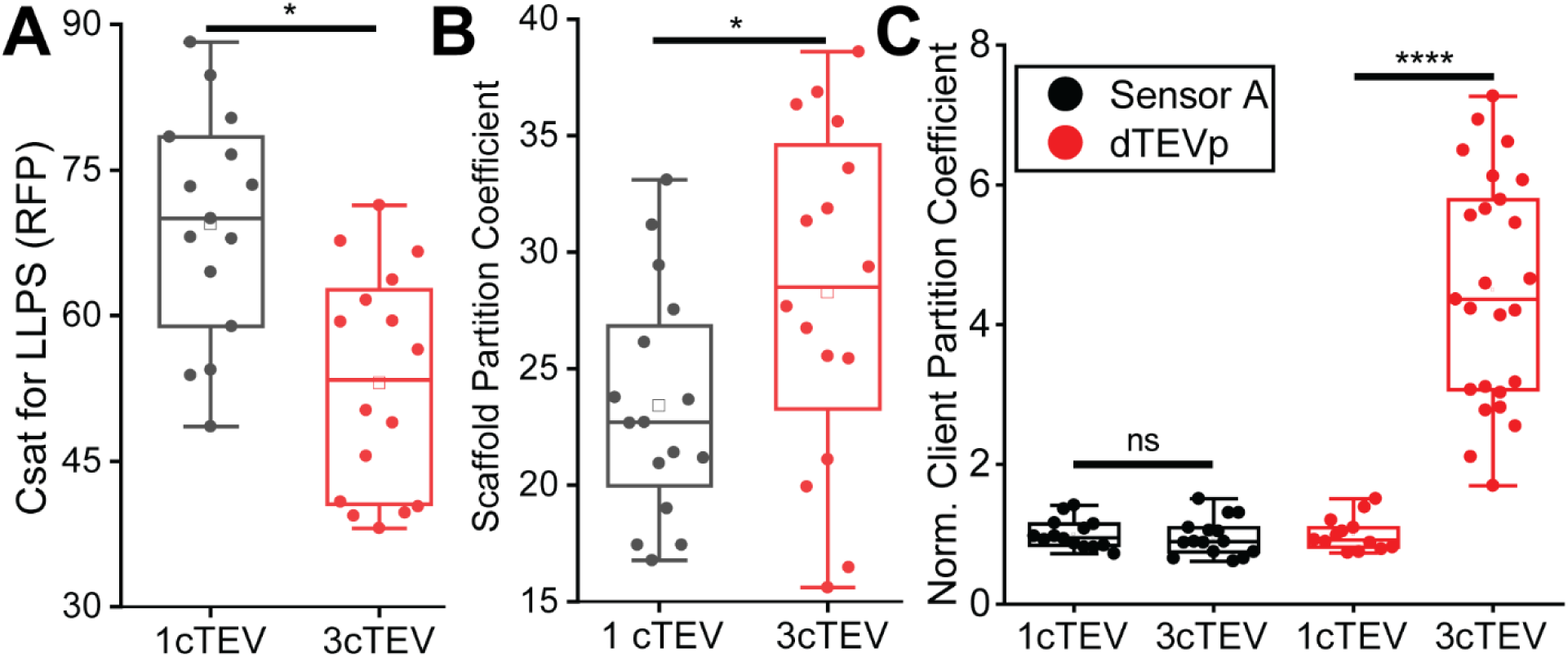
The number of client binding sites along a large IDP-scaffold alters its intracellular LLPS dynamics and client recruitment. **(A-B)** Scaffold critical concentration for LLPS **(A)** and partition coefficient **(B)** for 4-repeat r8 scaffolds that harbor either three (S1 in Fig. 6) or one (S2 in Fig 6) TEVp binding sites (cTEV) along their amino acid sequence. **(C)** Normalized partition coefficients for Sensor A or dTEVp in the presence of 4-repeat r8 scaffolds that harbor either one or three cTEV sites, using the mean partition coefficient of the corresponding 1cTEV constructs as reference for normalization. Asterisks denote statistical significance (* = P≤0.05; **** = P≤0.0001). ns: not statistically significant (p>0.05).

**Fig S7:**
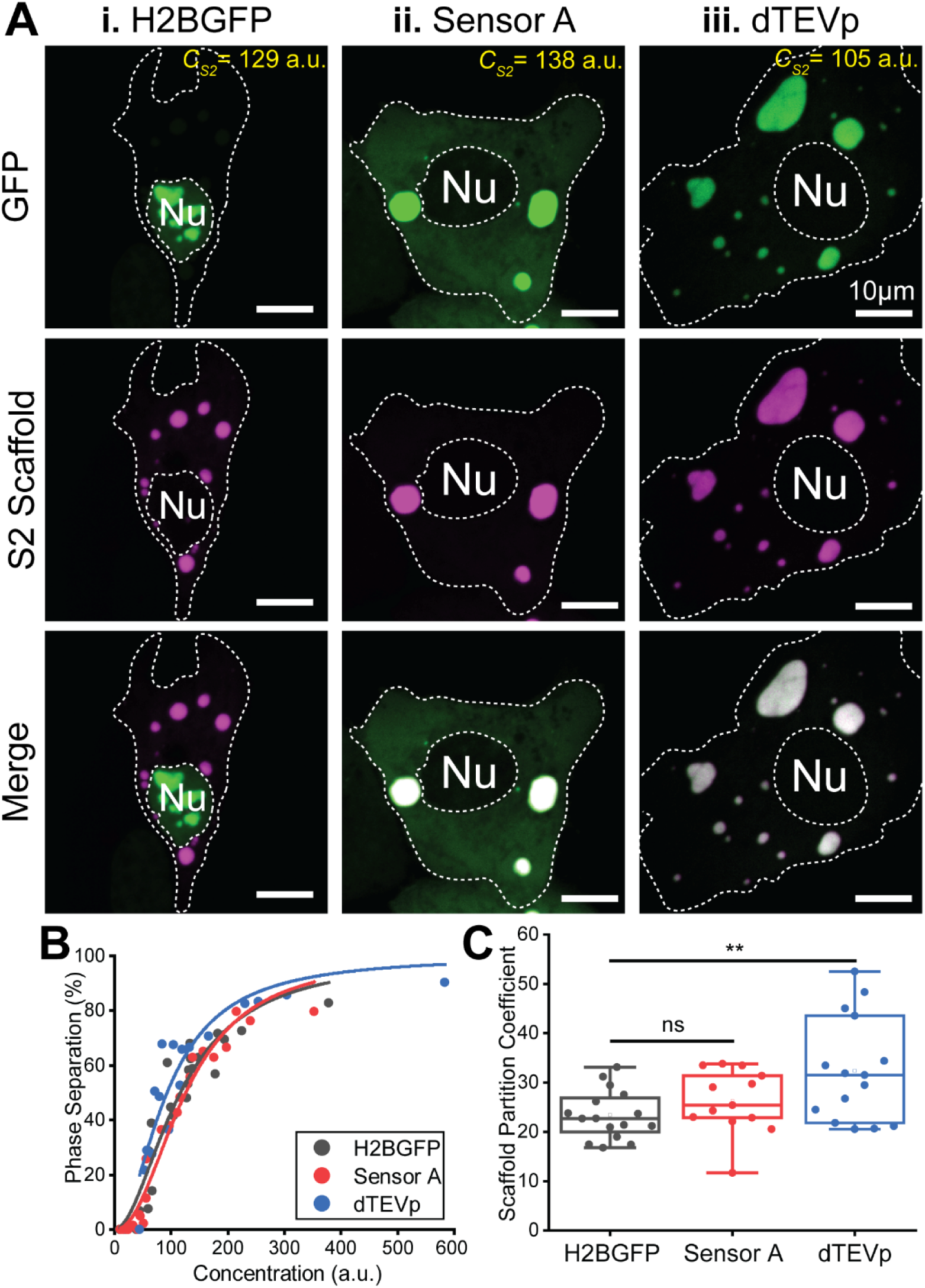
Intracellular LLPS behavior of large 4-repeat r8 scaffolds with one small binding site is sensitive to ligand-type client binding. **(A)** Live-cell images of (i) H2BGFP, (ii) Sensor A, (iii) dTEVp-client behavior in the presence of S2-scaffold (as in Fig. 6A) condensates. **(B)** Percent phase separation versus concentration of S2-scaffold in the presence of H2BGFP (grey), Sensor A (red), and dTEVp-client (blue). See Fig. 6G for quantifications of the dTEVp-induced shifts in the critical concentration for LLPS of the S2-scaffold. **(C)** Partition coefficients of S2-scaffold in the presence of H2BGFP (grey), Sensor A (red), and dTEVp-client (blue). Asterisks denote statistical significance (** = P≤0.01). ns: not statistically significant (p>0.05).

**Fig S8:**
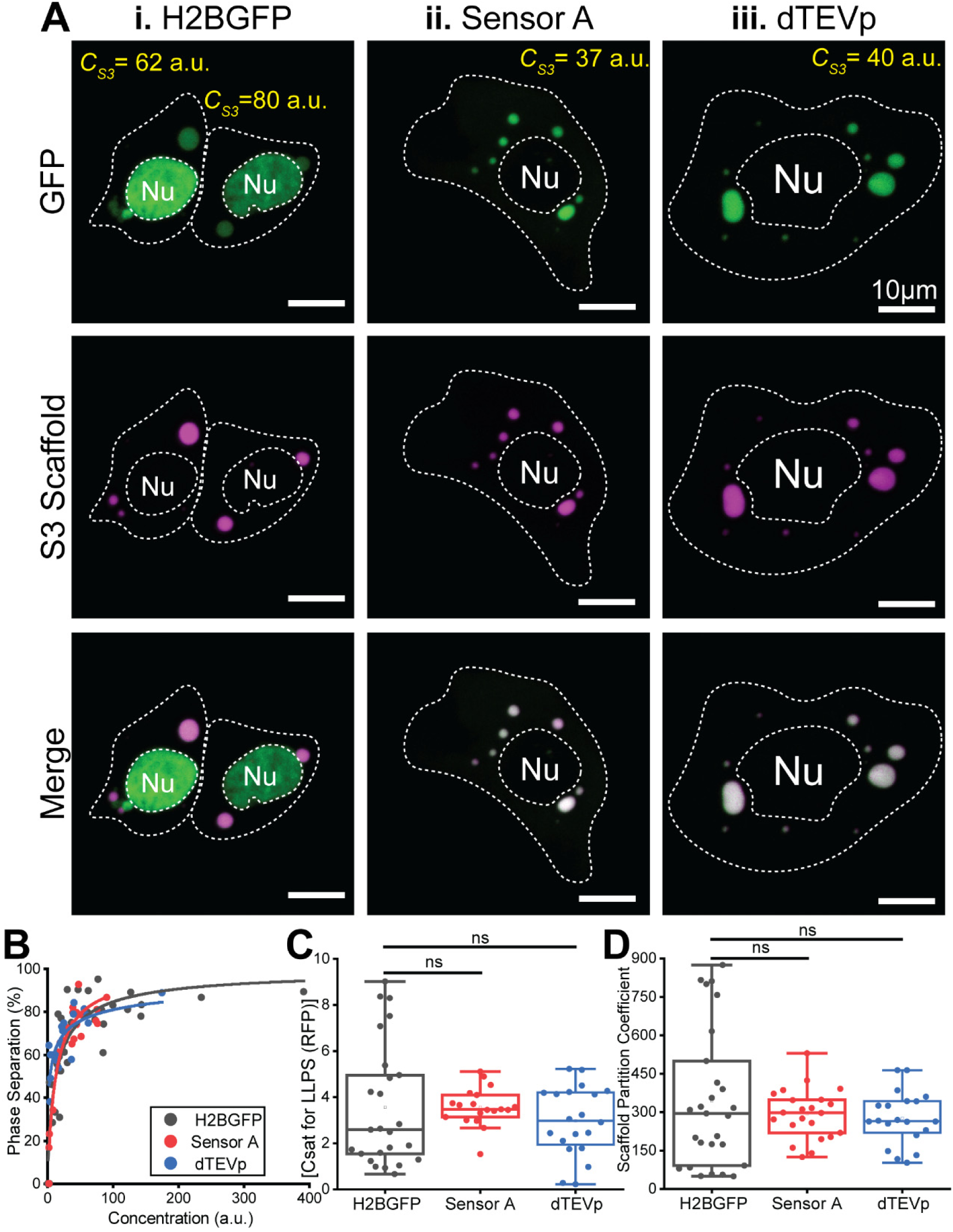
Intracellular LLPS behavior of very large 8-repeat r8 scaffolds is insensitive to ligand-type client binding. **(A)** Live-cell images of (i) H2BGFP, (ii) Sensor A, (iii) dTEVp-client behavior in the presence of S3-scaffold (as shown in Fig. 6A) condensates. **(B)** Percent phase separation versus concentration of S3-scaffold in the presence of H2BGFP (grey), Sensor A (red), and dTEVp-client (blue). **(C)** Critical concentrations for LLPS (C_sat_) of S3-scaffold in the presence of H2BGFP (grey), Sensor A (red), and dTEVp-client (blue). **(D)** Partition coefficients of S3-scaffold in the presence of H2BGFP (grey), Sensor A (red), and dTEVp-client (blue). ns: not statistically significant (p>0.05).

## Supplemental tables

**Table S1.**
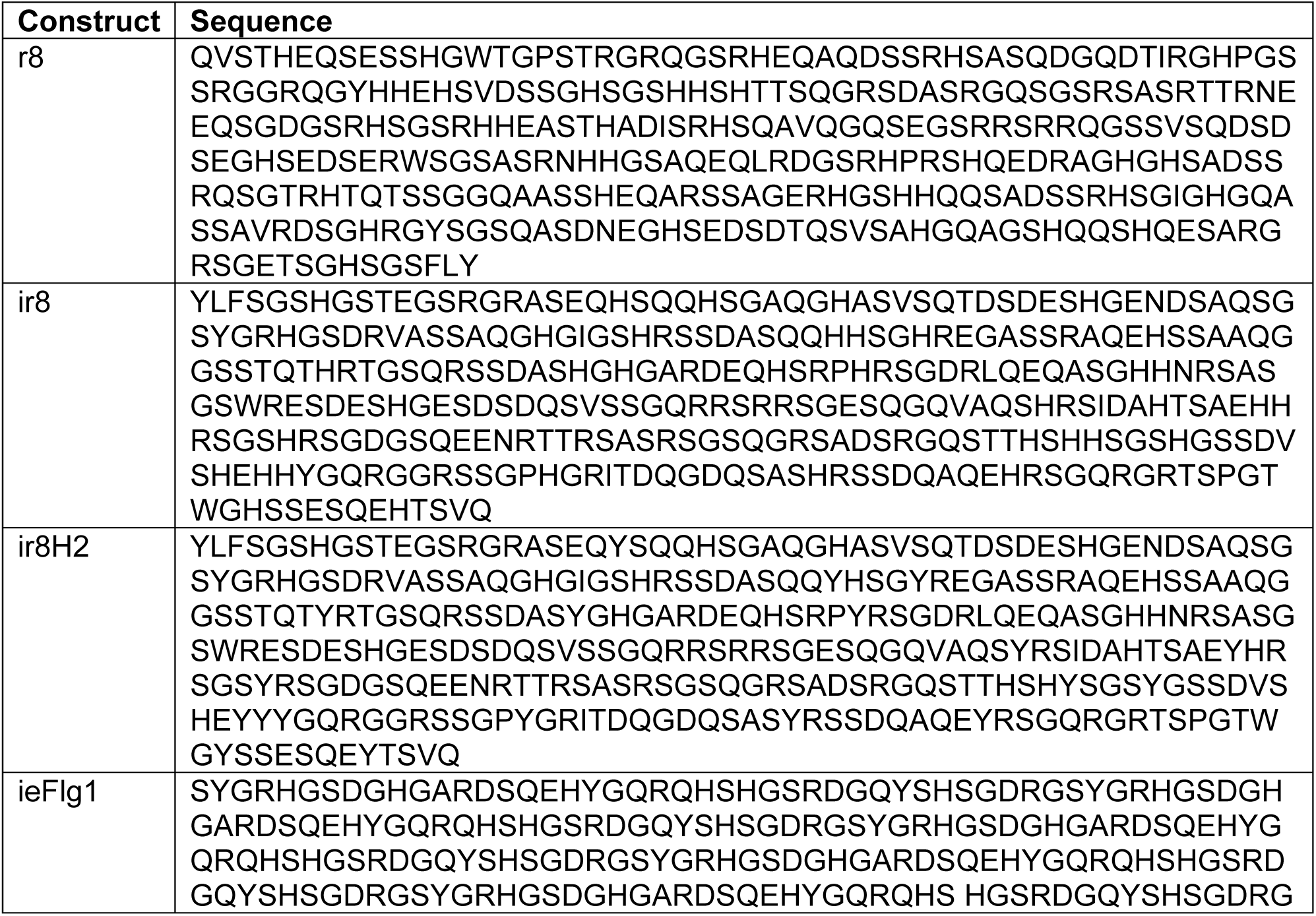
Sequence information for IDP-based sensing domains in Fig. 2 and Fig. S3.

**Table S2.**
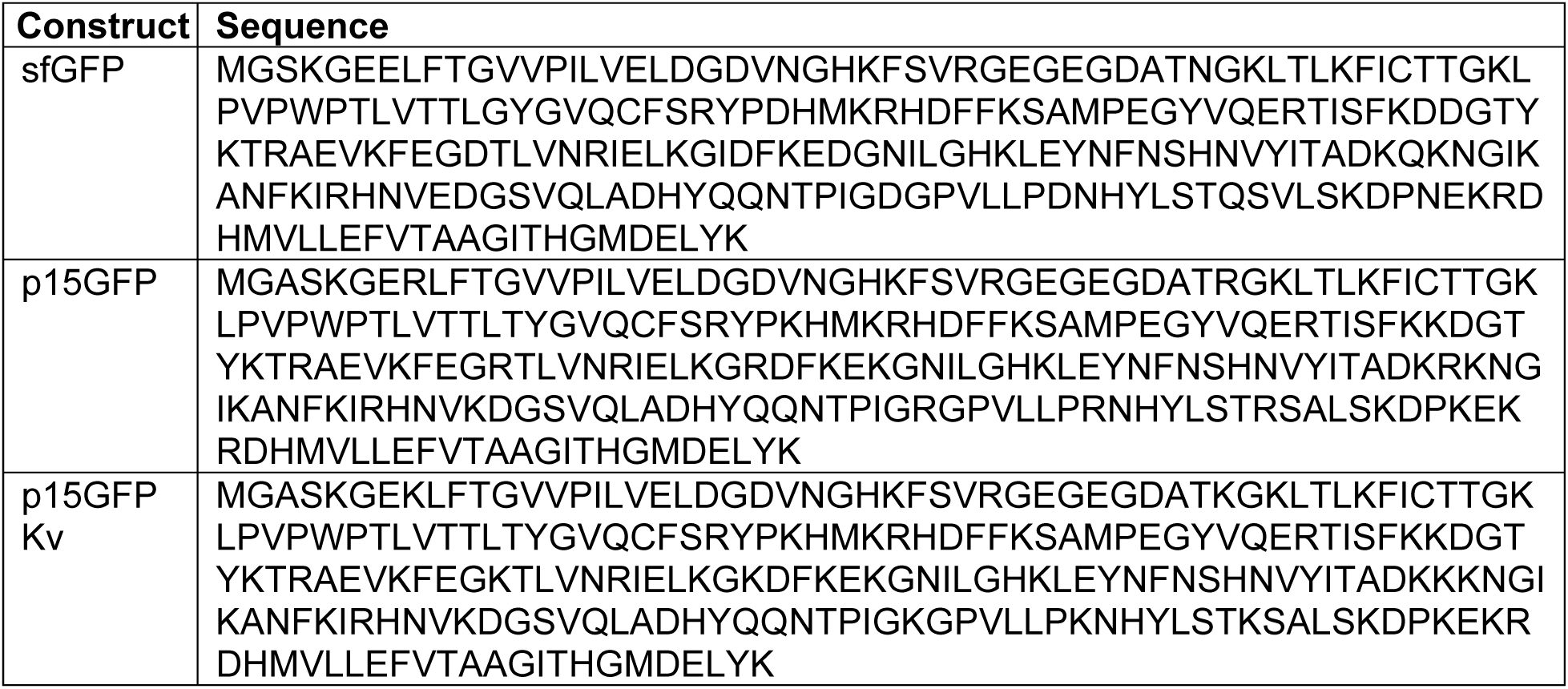
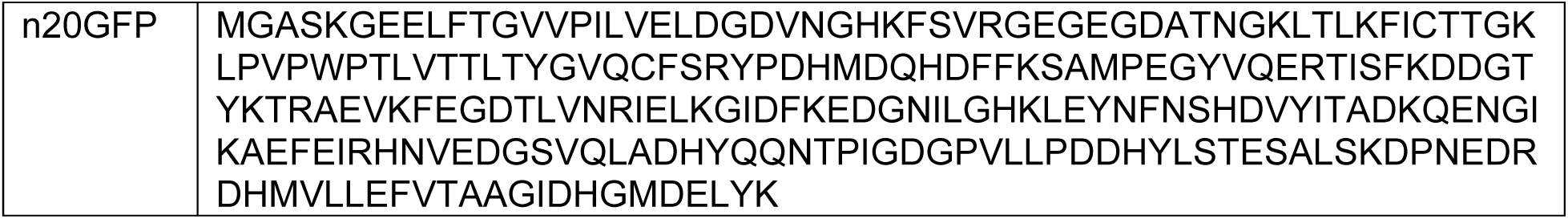
Sequence information for fluorescent proteins across sensor designs in Fig. 3.

**Table S3.**
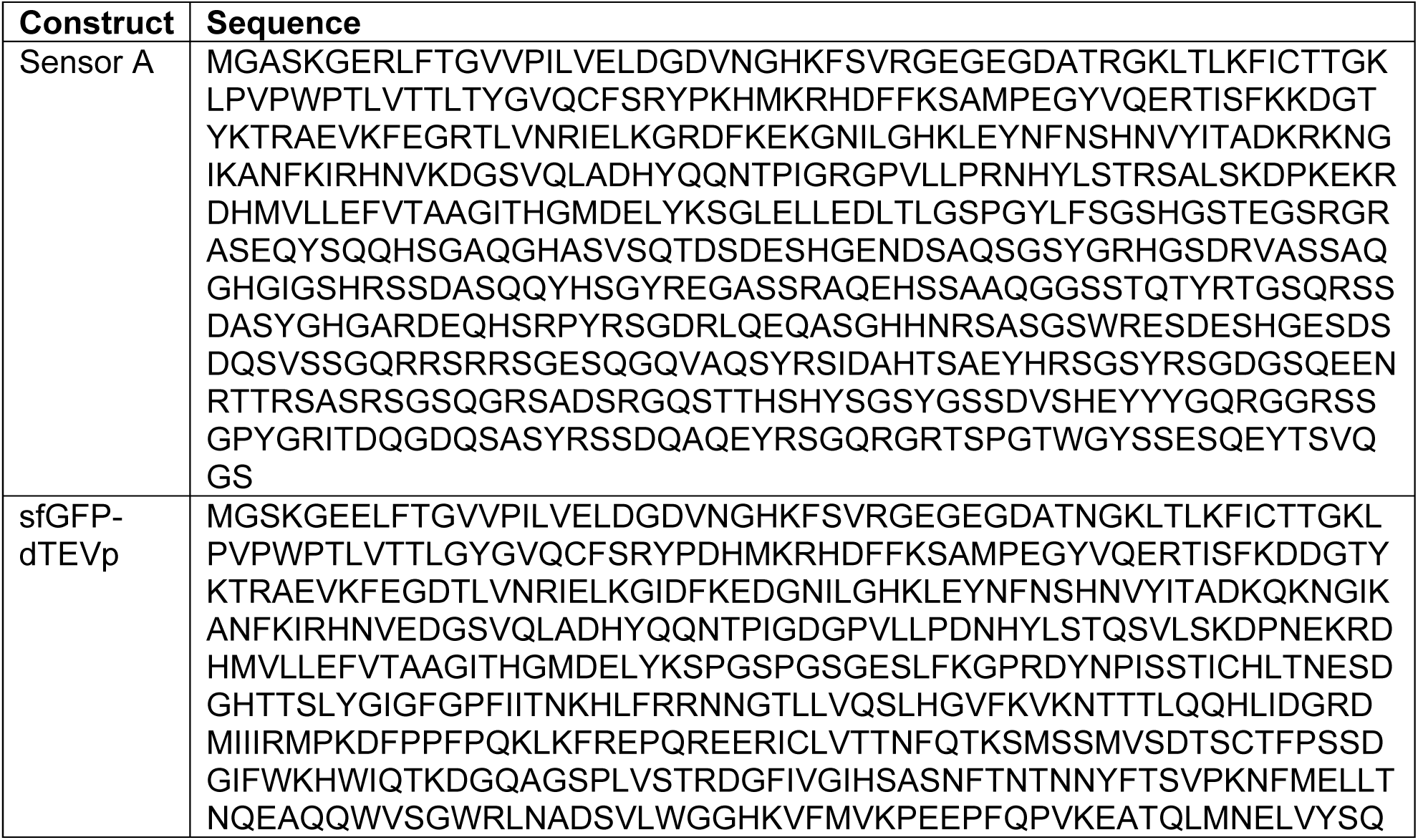
Sequence information for Sensor A and dTEVp-based client.

**Table S4.**
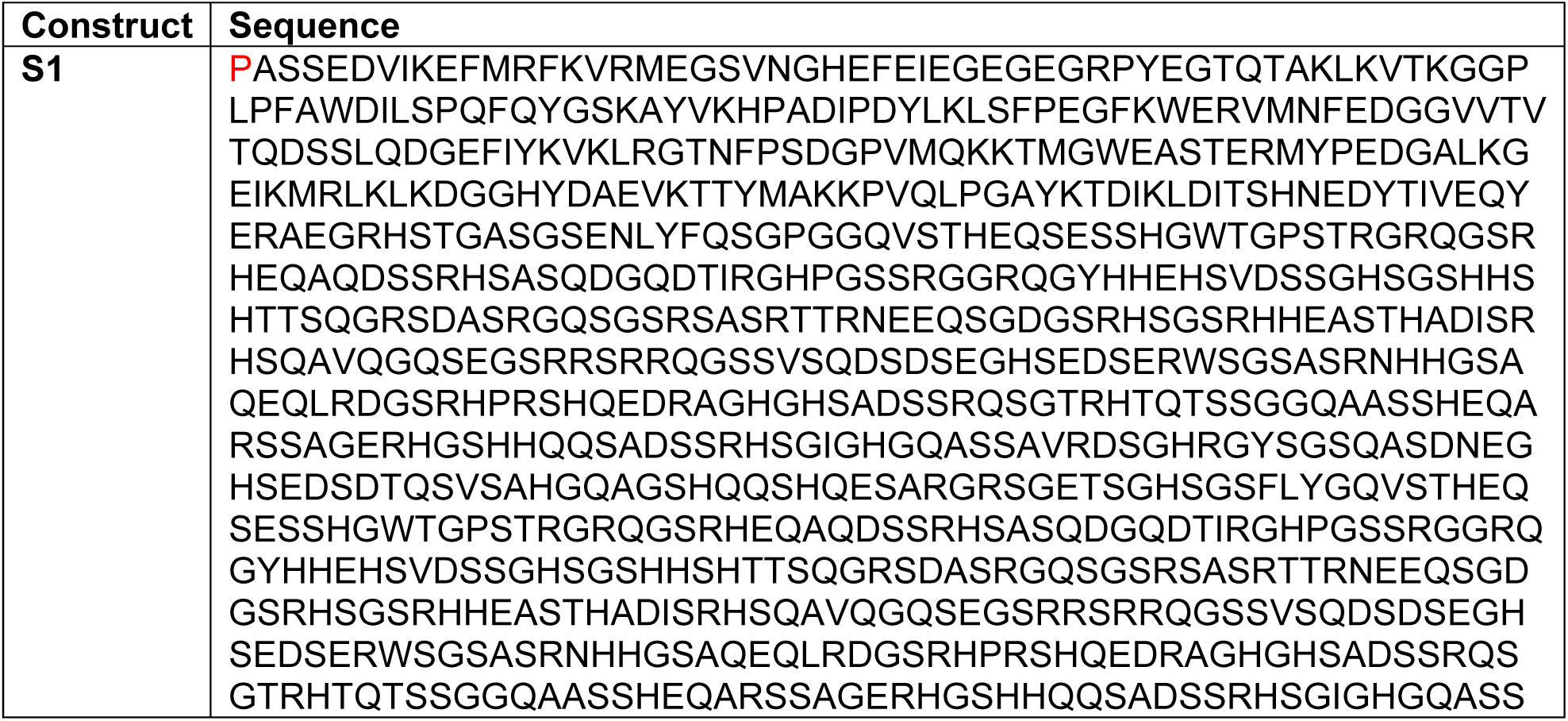
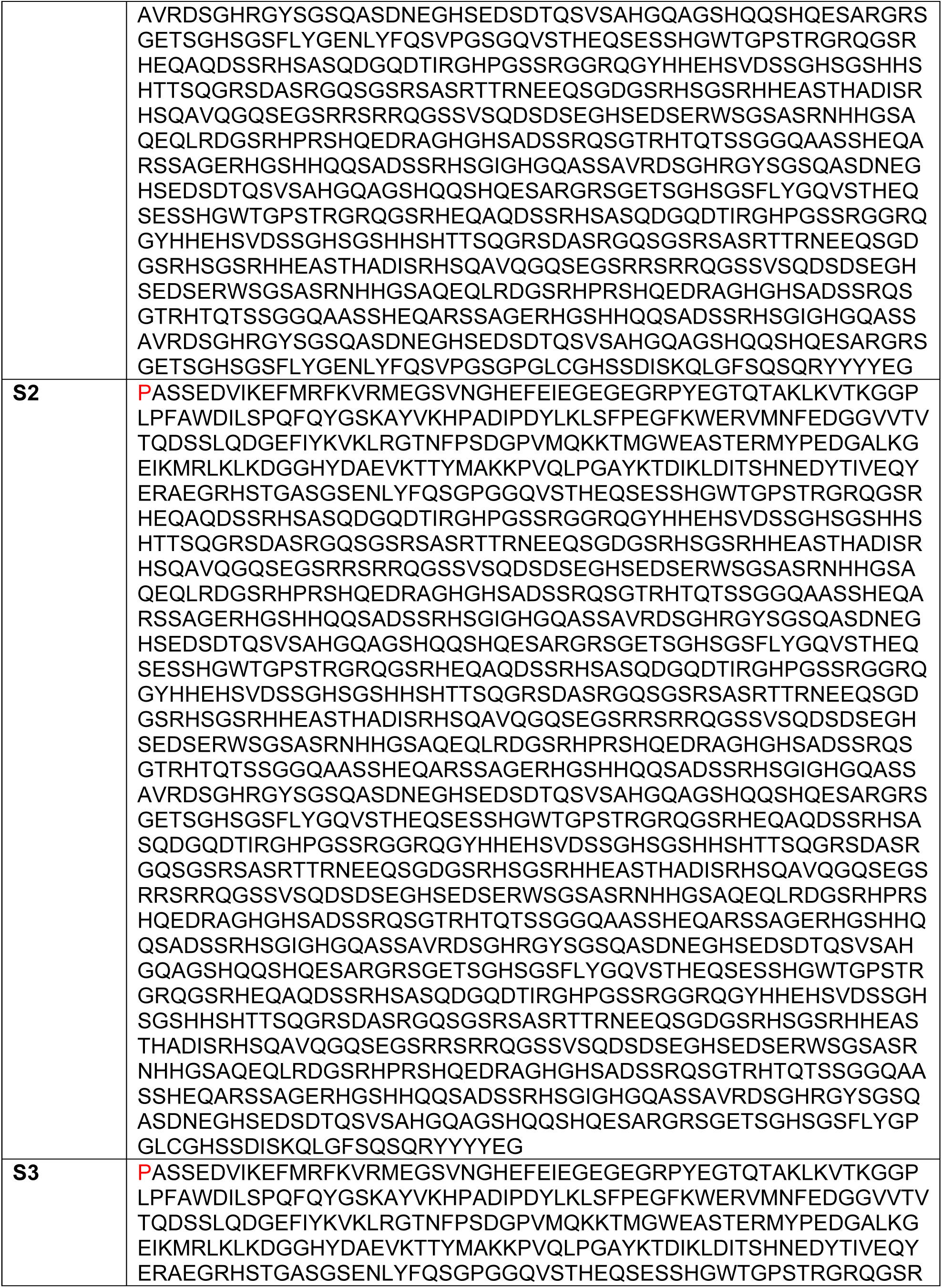
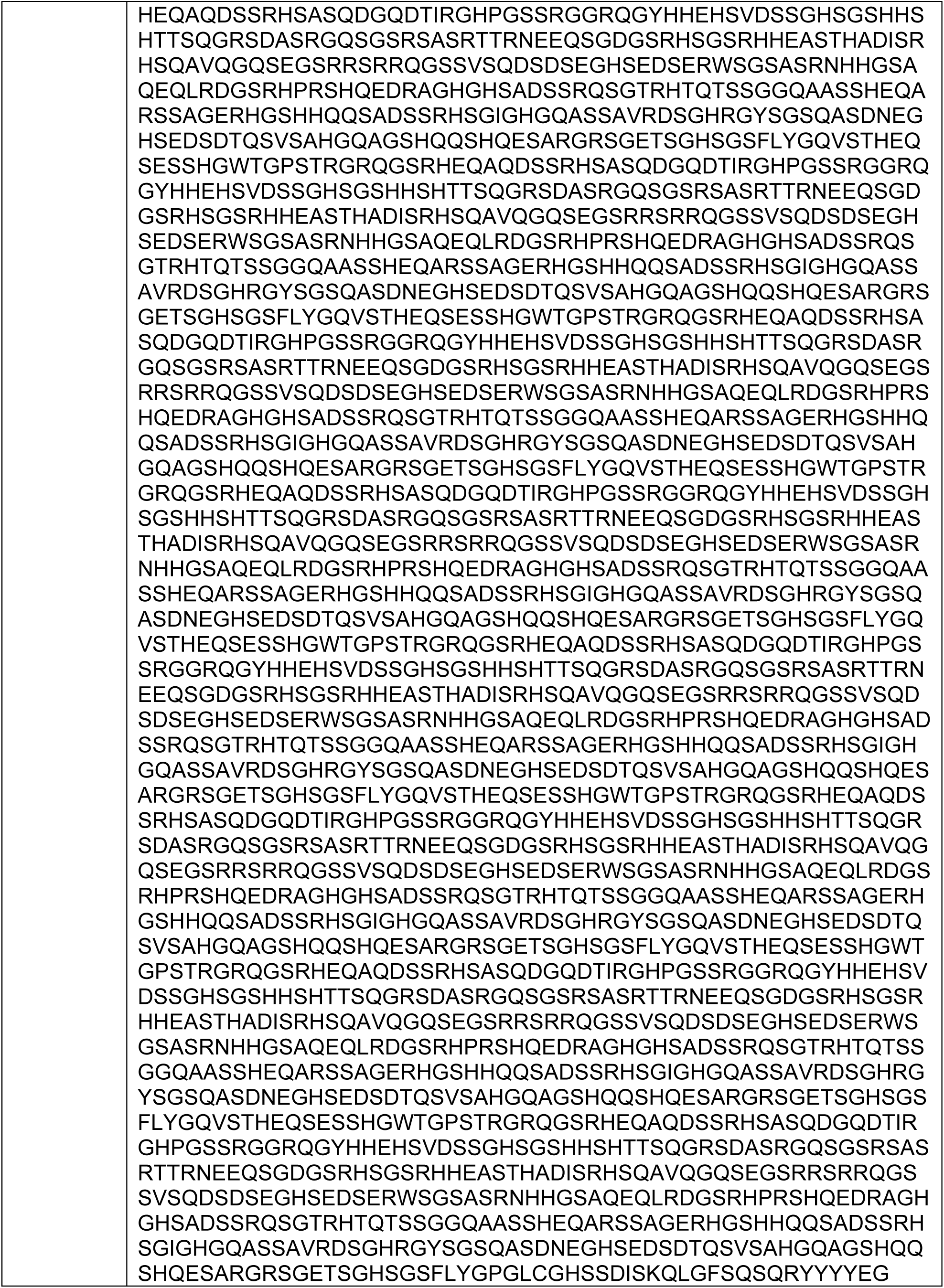
Sequence information for IDP-scaffolds (+Pro from P2A cleavage) in Fig. 6.

